# Spatially controlled microtubule nucleation and organization from crosslinker MAP65 condensates

**DOI:** 10.1101/2022.10.23.513406

**Authors:** Sumon Sahu, Prashali Chauhan, Ellie Lumen, Kelsey Moody, Karthik Peddireddy, Nandini Mani, Radhika Subramanian, Rae Robertson-Anderson, Aaron J Wolfe, Jennifer L. Ross

## Abstract

Microtubule organization in cells is essential for the internal structure and coordination of events of intracellular transport, mitosis, and cell motility. For many cell types, microtubule organization is dominated by centrosomal nucleation that use gamma-tubulin to template filaments. Yet, some cell types lack centrosomes or centrioles, such as plant cells. Instead, microtubules nucleate from regions with high concentrations of microtubule binding and nucleating proteins. A mechanism that can drive high local concentrations of nucleators is liquid-liquid phase separation of proteins with intrinsically disordered regions. Here, we report that the plant microtubule nucleator and crosslinking protein, MAP65-1, can form phase separated condensates at physiological salt and temperature without extra crowding agents. These condensates are liquid at first and can mature to gel-like phases over time and with different environmental conditions. We show that these condensates can nucleate and grow microtubule bundles that form asters, regardless of the viscoelasticity of the condensate. When gel-like droplets nucleate and grow asters from a shell of tubulin at the surface, the microtubules are able to re-fluidize the MAP65 condensate. Condensate-induced cytoskeletal formation could be a universal mechanism for organization of the microtubule and actin cytoskeletons in all cell types, especially cells without centrosomes.

## Introduction

Intracellular microtubule organization gives cells the structure and support to perform the myriad functions needed to coordinate their activities within tissues. Cell morphology, cell division, and even cell motility are driven by the microtubule cytoskeleton. Microtubule nucleation and growth must be regulated tightly in order to control the organization for each process in space and time. Microtubule-organizing centers (MTOCs) control nucleation and growth. In most cell types, the MTOC is synonymous with the centrosome containing the centriole, a complex made of gamma tubulin and other proteins organized into a barrel shape that can template microtubules directly. An important role of the centrosome is to organize the spindle poles in dividing cells (1). Yet, the centrosome is dispensable for even the most important processes, including mitosis (2, 3). For instance, the centrosome can be genetically eliminated (4–6) or ablated with a laser (7, 8), and the cell can continue through mitosis. Some cells, such as plant cells and meiotic animal egg cells, have no centrosomes (9, 10). In these cases, other organelles can nucleate microtubules in various regions of the cell including the perinuclear region (11–13), near chromosomes (14–16), from kinetochore fibers (17), and the Golgi complex (18, 19). Moreover, microtubules themselves can nucleate new microtubules from their surfaces to create branched networks (20, 21).

Recently, it has been shown that condensed, liquid droplets of microtubule-associated proteins (MAPs), such as tau, TPX2, CAMPSAP2, and BuGZ, Abl2, and +Tip proteins, can form condensates via LLPS and nucleate microtubules when free tubulin is added to these condensates (21–28). Tau is a good candidate for liquid-liquid phase separation (LLPS), since it is an intrinsically disordered protein (IDP) known to self-associate (29, 30), although it requires the addition of extra crowders to condense (22, 23). The ability of tau to form LLPS is affected by the sequence including known mutations that are observed in patients with dementia and other neurological disorders (31). The positive and negative effects of LLPS formation by tau are still being examined.

MAP65-1 (MAP65) is a microtubule-associated protein crosslinker from the PRC1/MAP65/Ase1 family that can promote nucleation of the microtubules (32). Here, we demonstrate a possible mechanism for microtubule nucleation and organization by MAP65 via LLPS. We show that both MAP65 and PRC1 can condense into liquid-like droplets in vitro without added crowding agents. Additionally, we show MAP65 droplets age into viscoelastic gels. When free tubulin is added, both the liquid and gel-like condensates concentrate tubulin, nucleate and grow microtubule in a spatially controlled manner to create microtubule organizations such as asters and spindle-like tactoid assemblies. Interestingly, we observe MAP65 mobility on microtubule aster bundles created from aged condensates is faster than aged condensates themselves, implying that the nucleated microtubules can deage MAP65 condensates, reversing hardening. Collectively, this work suggests that phase separated MAP65 condensates can act as a non-centrosomal nucleation and growth center for microtubules in plant cells.

## Results

### MAP65 undergoes liquid-liquid phase separation

In this study we used full length WT MAP65-1 (MAP65) from *Arabidopsis*, which is part of the Ase1/PRC1/MAP65 family. The full-length MAP65 has four rod and spectrin domains along the sequence and intrinsically disordered regions (IDRs) in the C-terminal tail (Fig. 1Ai) as predicted by two different PONDR algorithms (VSL3 and VL3, Fig. 1Aii) (33). Moreover, we find blocks of locally concentrated positive and negative charge regions, predicted by CIDER net charge per residue (NCPR), in the IDR regions (Fig. 1Aiii). These molecular properties are known markers of proteins that can condense, making MAP65 a good candidate to form condensates.

**Fig. 1.**
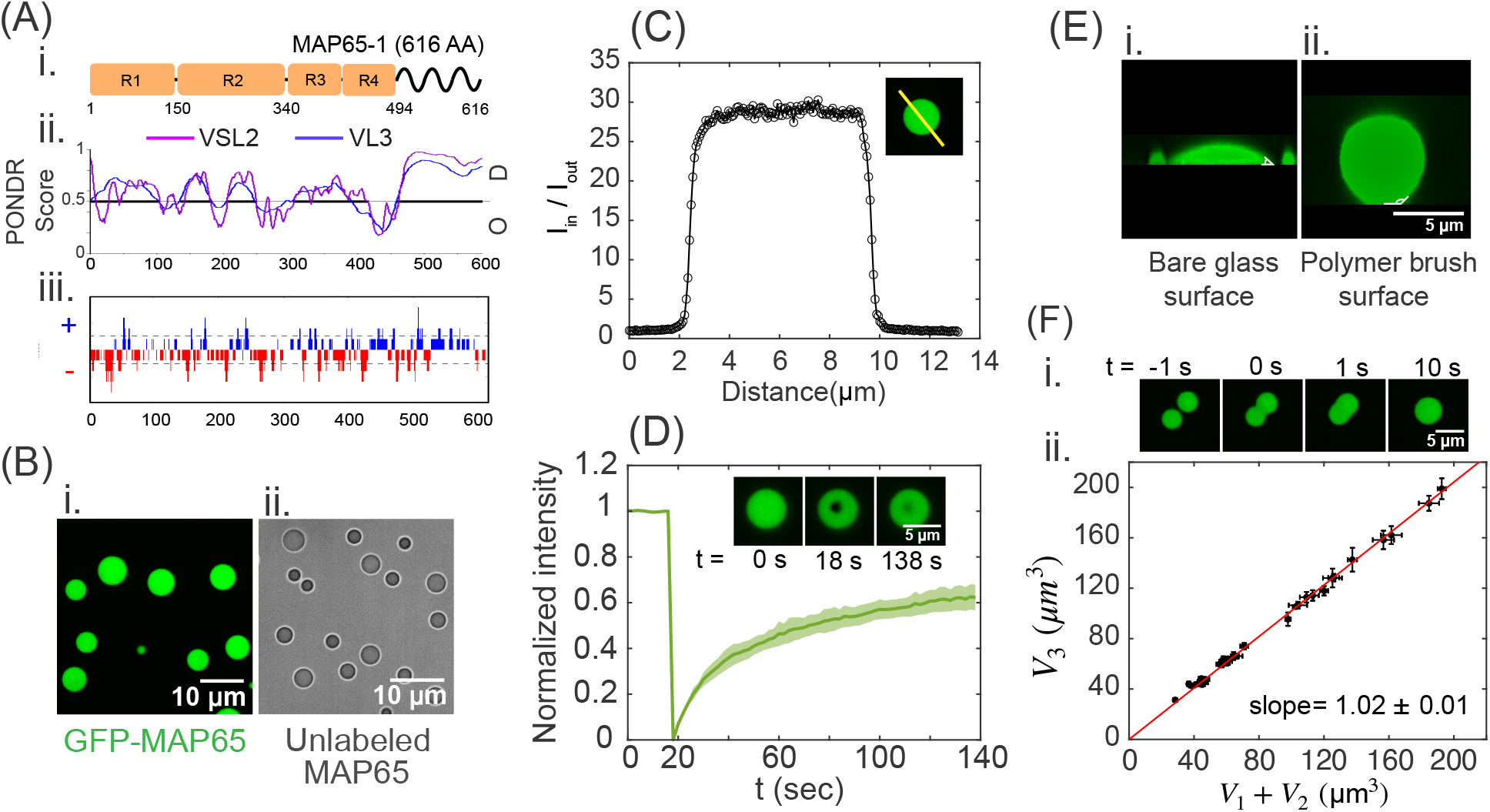
MAP65 can form liquid condensates. (A) (i) Cartoon of MAP65 sequence regions. *R*_1_, *R*_2_, *R*_3_, and *R*_4_ boxes represent spectrin domains. Tail domain is an IDR. (ii) Predicted disorder probability from PONDR algorithms, VSL2 (magenta) and VL3 (blue). Score > 0.5 is considered disordered. (iii) Predicted net charge per residue (NCPR) distribution from CIDER with positive charges (blue) and red charges (red). (B) Images of phase separated MAP65 in (i) confocal fluorescence or (ii) transmitted light microscopy. Scale bars are 10 *μm*. (C) Quantification of the partition coefficient, p, from the intensity profile of a confocal slice through the center of a droplet. Intensity is normalized so that the background level is one. (D) Quantification of FRAP normalized such that the initial intensity is one and the minimum intensity is zero with average (dark green line) and standard deviation (green shaded region) displayed (N=8). Image inset: FRAP sample data with droplet before bleach (t=0 sec), at bleach (t=18 sec), and after recovery (t=138 sec). Scale bar is 5 *μ*m. (E) Confocal images of droplets in the X-Z plane for MAP65 condensates (i) on bare glass, and (ii) on a polymer brush surface. Scale bars are 5 *μ*m. (F) (i) Images of droplet fusion over time where t = 0 sec indicates the frame where fusion started. (ii) Plot of the volume of the sum of the droplets’ volume before fusion, *V*_1_ + *V*_2_, and after fusion, *V_3_* (n=35). The best fit slope is 1.02 ± 0.01, indicating that volume is conserved.

We observed that purified, full-length MAP65 forms condensates via LLPS at 22 ±1 °*C* in a typical tubulin polymerization buffer, PEM-80 (80 mM PIPES, 1 mM EGTA, 2 mM MgCl_2_) at pH 6.8 (Fig. 1B). For confocal imaging, 10% of MAP65 was tagged with green fluorescent protein MAP65 (GFP-MAP65), which allowed us visualize round droplets using confocal fluorescence imaging (Fig. 1Bi). To ensure that GFP is not driving condensation, we imaged unlabeled full-length MAP65 using transmitted light microscopy in the same conditions and found that unlabeled MAP65 also formed condensates (Fig. 1Bii).

From confocal imaging, we observed that the MAP65 droplets are a single condensed phase that is composed of a MAP65-rich domain separated from the surrounding solution. There are no inhomogeneities or higher density regions within the droplet. We quantified the partition coefficient, p, to determine the relative difference in concentration between the droplet inside and outside using the intensity profile through a single confocal slice in the middle of the droplet, *p* = *I_in_/I_out_* (Fig. 1C), where *I_in_* and *I_out_* are the average intensity inside and outside the condensate, respectively. We find that *p* = 20.5 ±0.3 when the MAP65 concentration is 14 *μM* (mean ±SEM, N = 2466 droplets). Droplets can form when there is only 1 *μ*M MAP65 (SI text, Fig. S2). As the MAP65 concentration increases, the size, total number, and partition coefficient of droplets increases, but the partition coefficient saturates above 10 *μ*M (SI text, Fig. S2).

Using fluorescence recovery after photobleaching (FRAP), we test the mobility of the molecules within a condensate. We use a 405 nm laser to photobleach a small region within the droplet and image recovery (Fig. 1D). We quantify the fluorescence of the region over time, as described in the supplement (SI text, Fig. S1). The average fluorescence intensity of the spot rapidly recovers to 60% – 65% within a 2 min timescale (Fig. 1D). This data indicates that the MAP65 in the condensate is mobile and liquid-like.

MAP65 condensates are able to wet a glass cover slip surface when the glass is untreated. Using a confocal z-stack, we reconstruct and measure the contact angle for wetting the surface. For bare glass, the contact angle, *θ* ~ 50° (Fig. 1Ei). Wetting the surface inhibits further experiments such as fusion experiments to characterize condensate size and material properties. To reduce wetting, we pre-coat the cover glass surface with a polymer brush, Pluronic-F127, described in previous papers (34, 35). The polymer brush surface prevents protein adsorption and facilitates surface dewetting with an approximate contact angle *θ* ~ 130° (Fig. 1Eii). This surface coating allows MAP65 condensates to diffuse on the surface. When two MAP65 condensates come close and touch each other by thermal motion, they fuse into a single droplet rapidly (Fig. 1F).

Imaging confocal z-stacks over time during merging events, we quantify the droplet volumes before and after fusion. We repeated the experiment using transmitted light imaging to ensure there are no discrepancies due to using fluorescence. Plotting the volume of the merged final droplet, *V*_3_, as a function of the summed volume of the two droplets before merging, *V*_1_ + *V*_2_, we find that the slope is one, as expected if the volume is constant before and after droplet merging (Fig. 1Fii).

MAP65 is the plant analog of the PRC1 protein from mammalian cells that organizes microtubules during mitosis. Full-length PRC1 has similar domains as MAP65, including spectrin domains and an intrinsically disordered tail, similar charge distributions, and similar predictions for unstructured regions. We demonstrate that PRC1 can also form condensates (SI text, Fig. S3).

Many protein condensates are sensitive to environmental parameters such as temperature and ionic strength of the environment (36). The addition of salt inhibits MAP65 condensate formation, reducing both the condensate number density and droplet size (SI text, Fig. S4). The theoretical isoelectric point (pI) of MAP65 is 5.07 estimated using ExPasy Prot-param, suggesting that MAP65 is overall negatively charged at pH 6.8, but the charge distribution has positive and negative charges distributed along the length with a positively charged tail (SI text, Fig. S4). Unsurprisingly given their similar charge nature, both MAP65 and PRC1 show reduced condensate formation with increasing monovalent salt concentration (SI text, Fig. S4).

We find that temperature affects MAP65 condensation in an unexpected manner (SI text, Fig. S4). At low temperature 4 °*C*, many small droplets form. At room temeperature, the droplets were large and round, but at high temperature, 37 °*C*, droplets were non-spherical and adhered to one another indicating nucleation and growth of condensates with incomplete merging events between multiple droplets that are no longer liquid. This data implies that the temperature causes a transition from liquid to gel-like condensates, a process termed aging. For PRC1 at 37 °*C*, condensates are still round and liquid-like; they do not display aging (SI text, Fig. S4).

### Aging of condensates

Several proteins such as FUS, hnRNPA1, hnRNPA2, EWS, TAF15, and FIB1 are reported to form condensates that become gel-like with time (37–46). The change in the material properties from fluid-like to viscoelastic gels has been referred to as aging, hardening, or maturation of droplets. Aging can be probed using FRAP to quantify molecular mobility within the droplet and droplet fusion events to measure the surface tension changes (Fig. 2A). After maturation, mobility and merging cease. Droplets stick, but cannot coalesce because the molecules are jammed, as observed for MAP65 condensates at high temperature (SI text, Fig. S4).

**Fig. 2.**
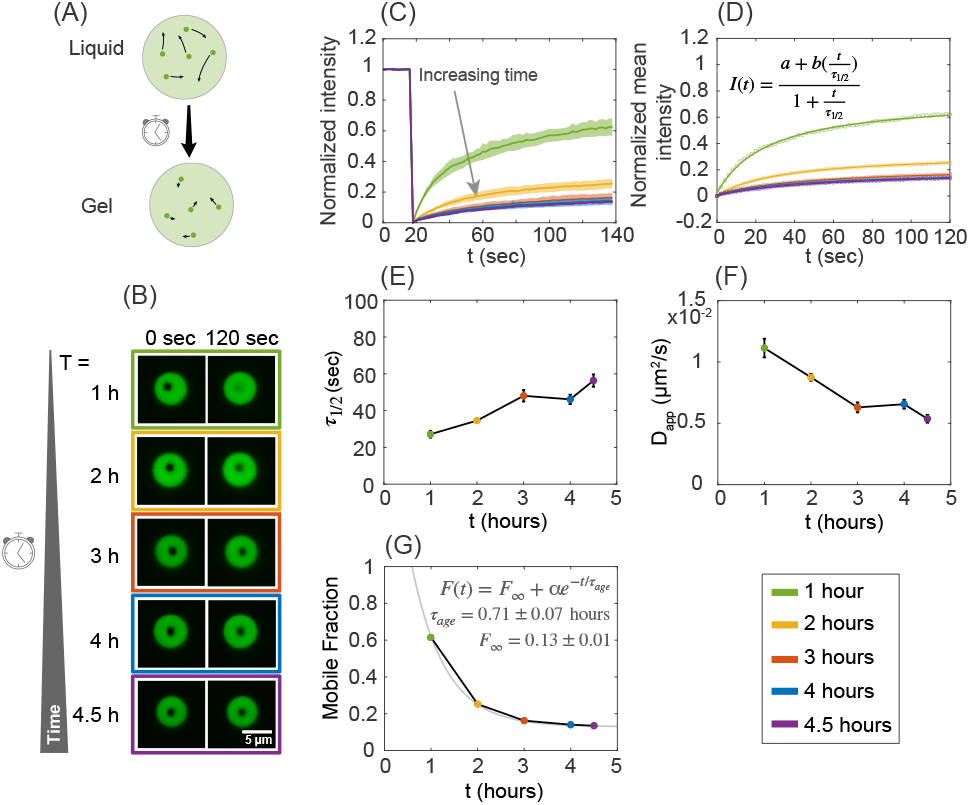
MAP65 droplet aging characterization using FRAP. (A) Cartoon schematic of a condensate transitioning from liquid-like to gel-like over time. (B) Time series images of droplet FRAP over maturation time. Left images indicate the initial photobleaching at t = 0 s. Right images show the same droplet 2 minutes later. The color of the box outline corresponds to the color of the data points in subsequent panels. The scale bar is 5 *μm*. (C) Normalized FRAP recovery curves for each five time points for maturation times of 1 hour (N=8, green), 2 hours (N=10, yellow), 3 hours (N=9, red), four hours (N=10, blue) and 4.5 hours (N=6, purple). The shaded regions indicates the standard deviation around the mean. (D) Normalized mean intensity recovery curves fit with the eqn. 1 to quantify recovery time constant *τ*_1/2_. (E) The recovery constant *τ*_1/2_ plotted against maturation time. The bars indicate a 95%confidence interval. (F) Apparent diffusion coefficient calculated from *τ*_1/2_ is plotted against maturation time. The bars indicate a 95%confidence interval. (G) Mobile fraction data shows exponential decay plotted against maturation time. The exponential decay fit (grey line) yields a characteristic aging time *τ_age_* = 0.71 ±0.07 hours with mobile fraction at long time, *F*_∞_, to be ~ 13%.

Here, we test the effects of time on MAP65 droplet maturation using FRAP to measure molecular mobility (Fig. 2B). Normalized fluorescence recovery curves are plotted and fit to this equation:

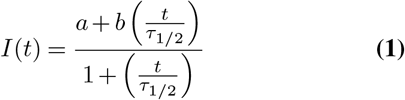

where *I*(*t*) is the intensity over time, *a* is initial intensity of the bleach, *b* is the intensity plateau at long times, and *τ*_1/2_ is the half-time of the recovery (Fig. 2D).

The *τ*_1/2_ values show an upward trend with the time that characterizes the reduced mobility of condensate aging. An hour after mixing, *τ*_1/2_ = 27.16 ±1.86 s, which jumps up two-fold at 4.5 hours, which is *τ*_1/2_ = 56.33 ±3.38 s (Fig. 2D,E, Table 1).

**Table 1.** MAP65 condensate aging data.

| Time (h) | $\tau_{1/2}$ (s) | $D_{app} \times 10^{-2} \mu\text{m}^2/\text{s}$ | Mobile fraction | Approx. Viscosity (Pa-s) |
| --- | --- | --- | --- | --- |
| 1 | $27.2 \pm 1.9$ | $1.11 \pm 0.07$ | 0.62 | 4.7 |
| 2 | $34.6 \pm 1.1$ | $0.88 \pm 0.03$ | 0.25 | 5.8 |
| 3 | $48.0 \pm 3.1$ | $0.63 \pm 0.04$ | 0.16 | 8.1 |
| 4 | $46.1 \pm 2.6$ | $0.66 \pm 0.04$ | 0.14 | 7.8 |
| 4.5 | $56.3 \pm 3.4$ | $0.54 \pm 0.03$ | 0.13 | 9.5 |

Using the mobility recovery time, we estimate an apparent diffusion coefficient, *D_app_*, for the MAP65 molecules inside the condensate: 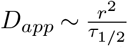, where *r* is radius of the photobleached region (47, 48), where *r* ~ 0.55*μ*m for our experiments (SI text, Fig. S1). The *D_app_* decreases as the droplets age over hours (Fig. 2F, Table 1). Using a second method (49), we find similar *D_app_* values, implying the method is robust.

To observe MAP65 molecule incorporation into the droplets from outside and mobility of MAP65 molecules at the droplet boundary, we also photobleached whole droplets at different maturation times (SI text, Fig. S5-S6). The mobility is quantified by plotting and fitting the radially averaged intensity profile from each droplet to the solution from 1D diffusion equation (SI text, Fig. S5-S6). We estimate the internal diffusion co-efficient, *D_in_* from the best fit (SI text, Fig. S6), and we find that the measured diffusion coefficient *D_in_* is similar to the apparent diffusion coefficient *D_app_* when using a small interior spot (Table 1, SI text, Table S1, Fig. S6).

The diffusion of protein molecules are coupled to the viscosity of the solvent by the Stokes-Einstein equation: 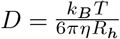, where *D* is the diffusion coefficient, *k_B_* is the Boltzmann’s constant, *T* is the temperature in *K*, *η* is the dynamic viscosity, and *R_h_* is the hydrodynamic radius of the protein. The hydrodynamic radius, *R_h_*, of MAP65, was reported to be ~ 42 Å (50). From *D* ~ *D_app_* and *R_h_*, we estimate the dynamic viscosity of the condensate as the droplets age (Table 1). At early times, the estimated viscosity is *η* ~ 4.7 Pa-s or 4670 mPa-s, which is similar to the viscosity of honey. For comparison, the dynamic viscosity of water at 22 °*C* is 0.95 mPa-s.

The rate of droplet merging can be quantified to find the surface tension of LLPS condensates (full method in SI text). When droplets are liquid-like, we plot time of droplet merging against the characteristic size of the merging droplets, and the slope is proportional to the ratio of viscosity to surface tension (SI text, Fig. S7 A). Using the viscosity found from *D_app_*, the surface tension of liquid-like droplets is *γ* ~ 12.3 *μN/m* in the first hour. At long times, droplets can no longer merge (SI text, Fig. S7 B), as expected for viscoelastic gels.

From the plateau value of the FRAP recovery curve, we estimate the mobile fraction of the molecules in the droplets. The mobile fraction dropped from ~ 60% after one hour of maturation, to ~ 14% after four and half hours. The reduction in mobile fraction was fit to an exponential form to reveal that a small fraction (~ 13%) of molecules remains mobile as the droplets age (Fig. 2G).

### Microtubule aster nucleation and growth from condensates

Condensates of microtubule-associated proteins can nucleate cytoskeletal networks by increasing the local concentration inside the condensate (21, 22, 24, 51). To test if MAP65 condensates can concentrate tubulin and nucleate microtubules, we perform experiments with tubulin added after droplets form. We examine the effect on microtubule organization as a function of droplet viscosity and size.

First, we examine the accumulation of tubulin into condensates by adding a low concentration of tubulin, [TUB] = 3.75*μM* without nucleotide to pre-formed MAP65 condensates made with 10*μM* MAP65. Droplets were examined 30 minutes after adding tubulin, and ~ 50% contained tubulin (Fig. 3A). An intensity scan of a confocal slice through the center of the droplet is used to determine the partition coefficient (Fig. 3Ai). We find that the average partition coefficient for tubulin inside compared to outside the droplet is *p* = 7.2±0.3 (mean ±SEM, N = 55 droplets). Given the total concentration of tubulin added, we estimate that the concentration inside these droplets is ~ 26 *μM* high enough to nucleate microtubules locally, if we had provided nucleotide to the system.

**Fig. 3.**
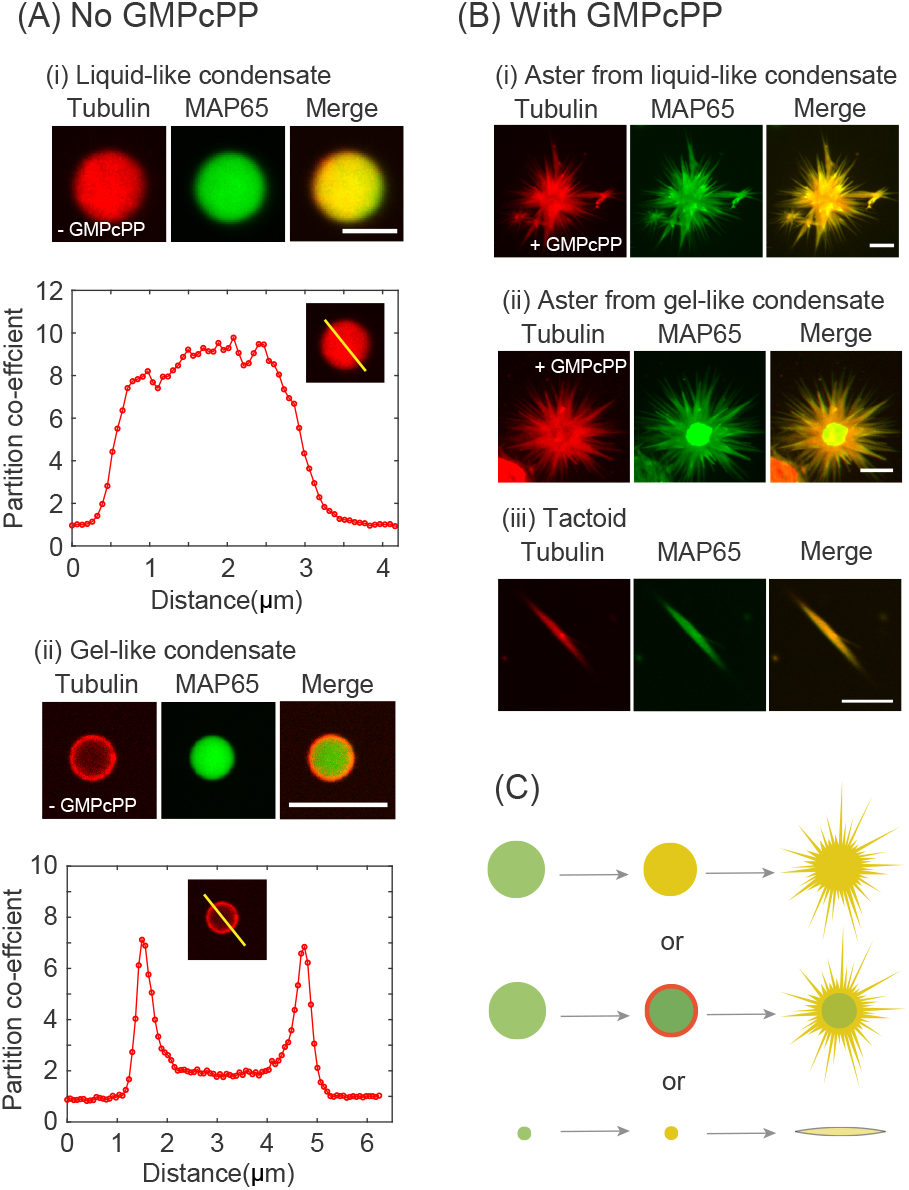
Tubulin colocalization and microtubule organization with MAP65 condensates. (A) Tubulin co-localizes with MAP65 liquid droplets. (i) Representative confocal image of rhodamine tubulin (red) and GFP-MAP65 (green) and overlay (merge) in a liquid-like droplet. Scale bar is 5 *μm*. Intensity scan through the droplet in the rhodamine channel normalized so that the intensity outside is one. (ii) Tubulin co-localizes with MAP65 aged droplets. Representative confocal images of rhodamine tubulin (red) and GFP-MAP65-1 (green) and overlay (merge) in a gel-like droplet. Scale bar is 5 *μm*. Intensity scan through the droplet in the rhoamdine channel normalized so that the intensity outside the droplet is one. (B) Microtubules nucleate from liquid-like or gel-like droplets. (i) Maximum projection of confocal z-stack shows that microtubule asters form when tubulin and GMPcPP is added to pre-formed MAP65-1 droplets that are liquid-like. (ii) Maximum projection of microtubule aster nucleated from gel-like MAP65 condensate. (iii) Microtubule tactoid, tapered bundle, nucleates from small MAP65 condensates. Scale bar is 5 *μm* for all images. (C) Cartoon schematic of mechanism of microtubule nucleation from MAP65 condensate showing that droplet viscosity and size result in different microtubule organizations.

When tubulin without nucleotide is added to MAP65 condensates that have aged and become more gel-like, tubulin is excluded from the MAP65 condensate core and forms a shell of tubulin on the condensate surface (Fig. 3Aii). The tubulin at the edge is highly enriched to *p* = 4.8 ±0.2 (mean ±SEM, N = 26 droplets). The intensity of tubulin inside the center of the droplet is higher than the background, but not as enriched, only 2.1 ±0.1 (mean ±SEM, N = 26 droplets) compared to the outside of the droplet. We estimate that the tubulin concentration in the ring is ~ 18 *μM*, which should be able to nucleate microtubules when there is nucleotide.

Next, we examine if the tubulin co-localization can nucleate and grow microtubules from the condensate. We add GMPcPP and the same tubulin concentration to pre-formed droplets at [MAP65] = 28*μM*. For liquid-like droplets, the tubulin is able to diffuse into the middle of the droplet and form microtubule asters with MAP65 bound to the microtubules (Fig. 3Bi). For gel-like MAP65 condensates, microtubule asters still form, but there is a core of MAP65 where tubulin is excluded at the center (Fig. 3Bii). MAP65 is found to distribute along the microtubules of the aster.

In addition to asters with many protruding microtubule bundles, we also observed microtubule tactoids in the same sample (Fig. 3Biii). Tactoids are finite sized, tapered bundles of microtubules which we have previously observed when microtubules were nucleated in the presence of MAP65 (34, 35). In this experiment, tactoids appear to be nucleated and grown from small MAP65 droplets, which likely only support bidirectional microtubule growth. It is possible that the finite size of the tactoids we previously observed were nucleated and grown from small MAP65 droplets that formed prior to tubulin nucleation and growth.

### Microtubules de-age gel-like condensates

To examine the mobility of each species in the condensates with tubulin, we perform two-color FRAP on gel-like MAP65 droplets that co-localized tubulin dimers or nucleated asters (Fig. 4A). In the absence of nucleotide, MAP65 recovery is minimal, with a smaller mobile fraction (2%) and similar recovery time compared to what was measured for gel-like MAP65 condensates without tubulin (Fig. 2). The tubulin FRAP shows an interesting biphasic recovery. We fit the normalized mean intensity with a double exponential equation, *I_TUB_*(*t*) = *A*(1 – *e*)^-*t/τ*_*TUB*, 1_^) + *B*(1 – *e*)^-*t/τ*_*TUB*, 2_^). The fast timescale is on the order of 5 s, and the slower recovery is 300 s (Fig. 4Aiii). We test if the center of the droplet is less mobile for tubulin by quantifying the recovery rates in the core compared to the edge of the droplet. We find the same biphasic recovery for the center of the condensate and the edge (data not shown), implying that tubulin can move with two distinct rates in any location within the MAP65 droplet. Next, we perform FRAP on microtubule asters formed around gel-like MAP65 condensates. We find that the mobile fraction of MAP65 is an order of magnitude higher and the recovery timescale is two times faster in the presence of the microtubules (Fig. 4B). The tubulin, which is now polymerized into microtubules, is less mobile in asters, and the recovery is still biphasic. The mobile fraction drops by a factor of three, the faster timescale increases by a factor of two, and the longer timescale increases by a factor of five (Fig. 4Biii). This result is surprising, since the aster is formed around an aged MAP65 condensate that has low MAP65 mobility without microtubules, indicating that the formation of the microtubules can release MAP65 from the gel-like core and alter the phase of the condensed MAP65. This phenomena is not a product of the tubulin, but requires microtubules to nucleate and grow.

**Fig. 4.**
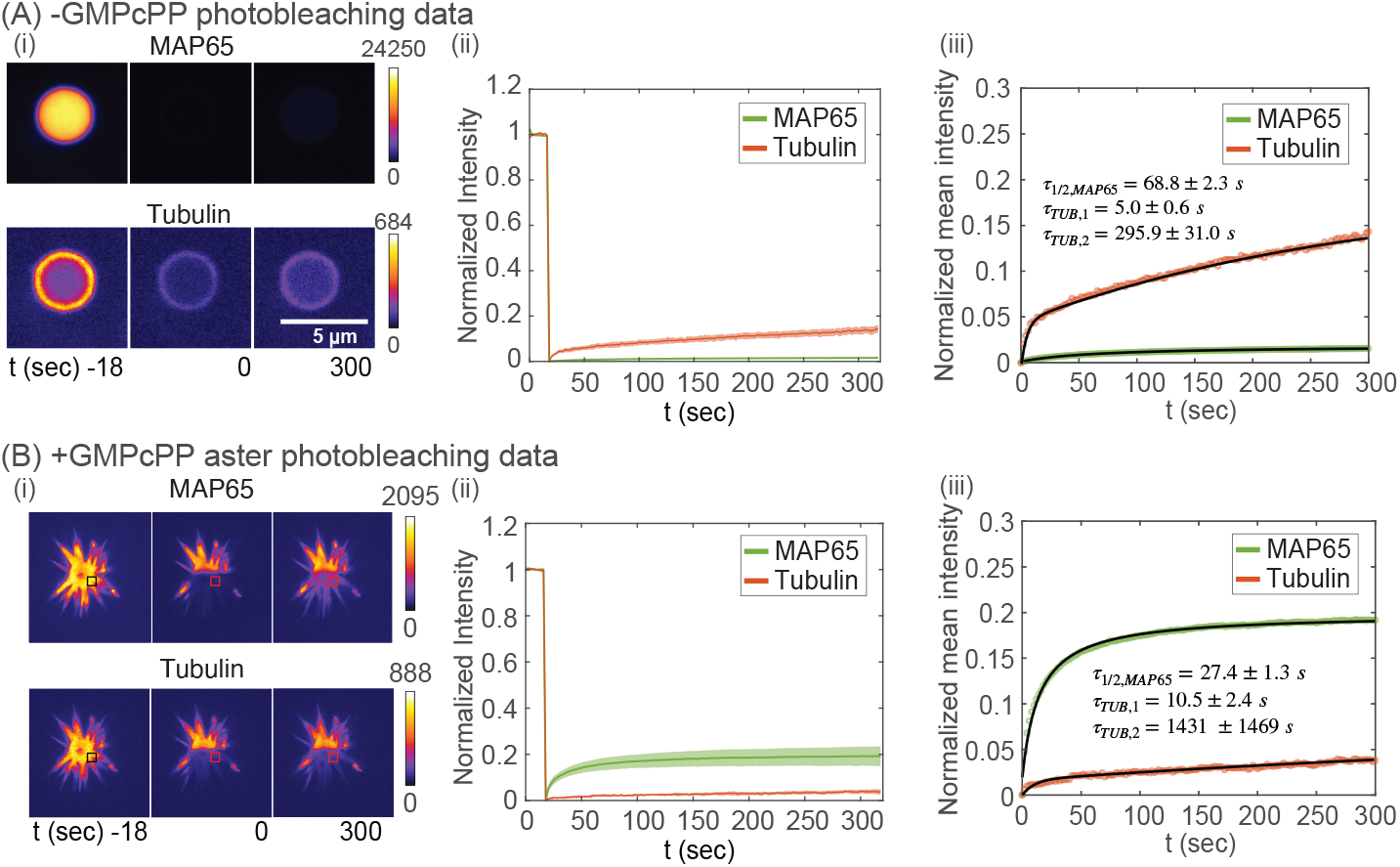
Photobleaching experiments on gel-like MAP65 condensates in presence of tubulin without and with nucleotide to form microtubules. (A) FRAP on tubulin. (i) Time series images of whole droplet FRAP for time points at t=−18 s, 0 s, and 300 s after photobleaching for GFP-MAP65 (top) and rhodamine-tubulin (bottom). Intensity portrayed using fire look up table, as denoted by color bar and ranges given. Scale bar is 5 *μ*M. (ii) Normalized FRAP recovery curved for GFP-MAP65 (green) and rhodamine-tubulin (red). The shaded region denotes the standard deviation around the mean (N=5). (iii) Normalized mean intensity curves fit with eqn.1 for GFP-MAP65 (green data, black line) or biphasic exponential recovery for tubulin (red data, black line). (B) FRAP on asters. (i) Time series images of whole droplet FRAP for time points at t=−18 s, 0 s, and 300 s after photobleaching for GFP-MAP65 (top) and rhodamine-tubulin (bottom). Intensity portrayed using fire look up table, as denoted by color bar and ranges given. Boxes represent area where analysis was performed. Scale bar is 5 *μ*M. (ii) Normalized FRAP recovery curved for GFP-MAP65 (green) and rhodamine-tubulin (red). The shaded region denotes the standard deviation around the mean (N=6). (iii) Normalized mean intensity curves fit with eqn. 1 for GFP-MAP65 (green data, black line) or biphasic exponential recovery for tubulin (red data, black line).

## Discussion

In this study we show that the microtubule-crosslinking proteins, MAP65 and PRC1 are both able to form condensates at low salt and moderate temperature in vitro (Fig.1, SI text, Figs. S2-S4). Excitingly, we find that this activity is temperature and salt-dependent, but does not require extra crowding agents, as observed for other MAPs, such as tau (22, 31), which is likely due to their molecular properties (Fig. 1, SI text, Fig. S3).

We find that MAP65 condensates age over time and transition from liquid-like to gel-like state. The gel-like state for MAP65 is never fully solid, since there is still a small mobile fraction (13%) that recovers even at 4.5 hours (Fig. 2). This implies that the core of the aged condensate is likely a viscoelastic network of MAP65 molecules, as opposed to a crystaline solid or a rigid aggregate.

Regardless of the maturation state of the MAP65 condensate, they are able to co-localize high concentrations of tubulin to cause localized nucleation and growth of microtubules (Fig. 3). This could allow MAP65 condensates to serve as microtubule-organizing centers in plant cells. Indeed, MAP65 family proteins are important for creating the phragmoplast in plant cells during mitosis (52, 53) and directing the cortical microtubule array in plants (32, 54).

The maturation of the droplet controls the localization of tubulin, creating a shell of tubulin when the droplet is aged (Fig. 3). Interestingly, the MAP65 mobile fraction in the presence of tubulin dimers is even smaller (2%), implying the tubulin shell impedes MAP65 mobility. Similar phase separations have been observed for traditional condensates from RNA-binding proteins and RNA with various viscoelasticity parameters for each phase (55). In our experiments, the tubulin shell appears to create a barrier for MAP65 diffusion.

One might expect that the barrier created by tubulin on aged MAP65 droplets could be exacerbated when the tubulin nucleates and grows microtubules, creating a cage around the MAP65 condensate. Instead of creating a larger barrier, we find that microtubules fluidize the gel-like MAP65 condensates - increasing both the mobile fraction and mobility of MAP65 molecules in the aged condensates (Fig. 4). Thus, the microtubules are able to reverse the gel-like phase change of MAP65 droplets, refluidizing and effectively “de-aging” the droplets.

The ability of microtubules to reverse the aging of droplets corroborates the idea that the mature condensates are viscoelastic networks. In the aged condensates, MAP65 molecules likely bind to each other to form a polymer network inside the less mobile core. We postulate that when microtubules form, they reduce the MAP65-MAP65 interactions in favor of MAP65-microtubule interactions. Further, although microtubules appear excluded from the condensate core, the microtubules at the edge are likely able to draw out and exchange MAP65 at the condensate edge, increasing the mobile fraction and decreasing recovery time (Fig. 4).

Microtubule organization depends on the viscosity and the size of the MAP65 condensates. Large droplets form asters, and small droplets form finite-sized microtubule tactoid bundles that were tapered at the ends. Such spindle-like microtubule assemblies are like liquid crystal tactoids which were previously observed for microtubules (34, 35) and actin (56, 57). A recent report demonstrated that condensates made of actin-bundling protein VASP can nucleate actin to make tactoid-shaped bundles (58).

The use of cytoskeletal-associated proteins to form condensates that can direct the organization of microtubules and actin is likely a general organizational principle in cells. Indeed, cytoskeletal fiber formation has recently been reported for numerous cellular and in vitro studies for microtubules (21, 22, 25) and actin (58–61). This may suggest that cells use this universal strategy to control cytoskeletal filament organization (62), but it is especially important for cell-types or regions of the cell that do not have centrioles to nucleate and grow microtubules in the traditional manner. Further, demonstrating that the aging of these condensates is not a barrier to nucleation and growth makes the condensate-based cytoskeletal organzation mechanism more useful and applicable.

## Materials and Methods

The detailed materials and methods are available in the SI Appendix.

## ACKNOWLEDGEMENTS

This work was funded by a grant from the National Science Foundation NSF BIO-2134215 to JLR and partially funded from the Keck Foundation to Dr. R. Robertson-Anderson, M. Das, M. Rust, and JLR.

## Bibliography

1. Mónica Bettencourt-Dias and David M. Glover. Centrosome biogenesis and function: Centrosomics brings new understanding. Nature Reviews Molecular Cell Biology, 8(6):451–463, 2007. ISSN 14710072. doi: 10.1038/nrm2180.

2. Alexey Khodjakov, Richard W. Cole, Berl R. Oakley, and Conly L. Rieder. Centrosome-independent mitotic spindle formation in vertebrates. Current Biology, 10(2):59–67, 2000. ISSN 09609822. doi: 10.1016/S0960-9822(99)00276-6.

3. Mónica Bettencourt-Dias. Q&A: Who needs a centrosome? BMC Biology, 11:1–7, 2013. ISSN 17417007. doi: 10.1186/1741-7007-11-28.

4. Greenfield Sluder. Two-way traffic: centrosomes and the cell cycle. Nature Reviews Molecular Cell Biology, 6(9):743–748, 2005. ISSN 1471-0080. doi: 10.1038/nrm1712.

5. Stephen Doxsey, Dannel McCollum, and William Theurkauf. Centrosomes in cellular regulation. Annual Review of Cell and Developmental Biology, 21:411–434, 2005. ISSN 10810706. doi: 10.1146/annurev.cellbio.21.122303.120418.

6. Vlastimil Srsen, Nicole Gnadt, Alexander Dammermann, and Andreas Merdes. Inhibition of centrosome protein assembly leads to p53-dependent exit from the cell cycle. Journal of Cell Biology, 174(5):625–630, 2006. ISSN 00219525. doi: 10.1083/jcb.200606051.

7. Jacob Odell, Vitali Sikirzhytski, Irina Tikhonenko, Sonila Cobani, Alexey Khodjakov, and Michael Koonce. Force balances between interphase centrosomes as revealed by laser ablation. Molecular Biology of the Cell, 30(14):1705–1715, 2019. ISSN 19394586. doi: 10.1091/mbc.E19-01-0034.

8. Sabrina La Terra, Christopher N. English, Polla Hergert, Bruce F. McEwen, Greenfield Sluder, and Alexey Khodjakov. The de novo centriole assembly pathway in HeLa cells: Cell cycle progression and centriole assembly/maturation. Journal of Cell Biology, 168(5): 713–722, 2005. ISSN 00219525. doi: 10.1083/jcb.200411126.

9. Han Zhang and R. Kelly Dawe. Mechanisms of plant spindle formation. Chromosome Research, 19(3):335–344, 2011. ISSN 09673849. doi: 10.1007/s10577-011-9190-y.

10. Julien Dumont and Arshad Desai. Acentrosomal spindle assembly and chromosome segregation during oocyte meiosis. Trends in Cell Biology, 22(5):241–249, 2012. ISSN 09628924. doi: 10.1016/j.tcb.2012.02.007.

11. Takahiro Hamada. Microtubule organization and microtubule-associated proteins in plant cells, volume 312. Elsevier Inc., 1 edition, 2014. ISBN 9780128001783. doi: 10.1016/B978-0-12-800178-3.00001-4.

12. Yiming Zheng, Rebecca A Buchwalter, Chunfeng Zheng, Elise M Wight, Jieyan V Chen, and Timothy L Megraw. A perinuclear microtubule-organizing centre controls nuclear positioning and basement membrane secretion. Nature Cell Biology, 22(3):297–309, 2020. ISSN 1476-4679. doi: 10.1038/s41556-020-0470-7.

13. Chris Ambrose and Geoffrey O. Wasteneys. Microtubule initiation from the nuclear surface controls cortical microtubule growth polarity and orientation in Arabidopsis thaliana. Plant and Cell Physiology, 55(9):1636–1645, 2014. ISSN 14719053. doi: 10.1093/pcp/pcu094.

14. Sylvain Meunier and Isabelle Vernos. Acentrosomal Microtubule Assembly in Mitosis: The Where, When, and How. Trends in Cell Biology, 26(2):80–87, 2016. ISSN 18793088. doi: 10.1016/j.tcb.2015.09.001.

15. Chaitanya A Athale, Ana Dinarina, Maria Mora-Coral, Céline Pugieux, Francois Nedelec, and Eric Karsenti. Regulation of Microtubule Dynamics by Reaction Cascades Around Chromosomes. Science, 322(5905):1243–1247, 2008. doi: 10.1126/science.1161820.

16. Maïwen Caudron, Gertrude Bunt, Philippe Bastiaens, and Eric Karsenti. Cell Biology: Spatial coordination of spindle assembly by chromosome-mediated signaling gradients. Science, 309(5739):1373–1376, 2005. ISSN 00368075. doi: 10.1126/science.1115964.

17. Helder Maiato, Conly L. Rieder, and Alexey Khodjakov. Kinetochore-driven formation of kinetochore fibers contributes to spindle assembly during animal mitosis. Journal of Cell Biology, 167(5):831–840, 2004. ISSN 00219525. doi: 10.1083/jcb.200407090.

18. K. Chabin-Brion, J. Marceiller, F. Perez, C. Settegrana, A. Drechou, G. Durand, and C. Poüs. The Golgi complex is a microtubule-organizing organelle. Molecular Biology of the Cell, 12 (7):2047–2060, 2001. ISSN 10591524. doi: 10.1091/mbc.12.7.2047.

19. Anna A.W.M. Sanders and Irina Kaverina. Nucleation and dynamics of Golgi-derived microtubules. Frontiers in Neuroscience, 9(NOV):1–7, 2015. ISSN 1662453X. doi: 10.3389/fnins.2015.00431.

20. Sabine Petry, Aaron C. Groen, Keisuke Ishihara, Timothy J. Mitchison, and Ronald D. Vale. Branching microtubule nucleation in xenopus egg extracts mediated by augmin and TPX2. Cell, 152(4):768–777, 2013. ISSN 10974172. doi: 10.1016/j.cell.2012.12.044.

21. Matthew R. King and Sabine Petry. Phase separation of TPX2 enhances and spatially coordinates microtubule nucleation. Nature Communications, 11(1):1–13, 2020. ISSN 20411723. doi: 10.1038/s41467-019-14087-0.

22. Amayra Hernández-Vega, Marcus Braun, Lara Scharrel, Marcus Jahnel, Susanne Wegmann, Bradley T. Hyman, Simon Alberti, Stefan Diez, and Anthony A. Hyman. Local Nucleation of Microtubule Bundles through Tubulin Concentration into a Condensed Tau Phase. Cell Reports, 20(10):2304–2312, 2017. ISSN 22111247. doi: 10.1016/j.celrep.2017.08.042.

23. Susmitha Ambadipudi, Jacek Biernat, Dietmar Riedel, Eckhard Mandelkow, and Markus Zweckstetter. Liquid-liquid phase separation of the microtubule-binding repeats of the Alzheimer-related protein Tau. Nature Communications, 8(1):1–13, 2017. ISSN 20411723. doi: 10.1038/s41467-017-00480-0.

24. Tsuyoshi Imasaki, Satoshi Kikkawa, Shinsuke Niwa, Yumiko Saijo-Hamano, Hideki Shigematsu, Kazuhiro Aoyama, Kaoru Mitsuoka, Takahiro Shimizu, Mari Aoki, Ayako Sakamoto, Yuri Tomabechi, Naoki Sakai, Mikako Shirouzu, Shinya Taguchi, Yosuke Yamagishi, Tomiyoshi Setsu, Yoshiaki Sakihama, Eriko Nitta, Masatoshi Takeichi, and Ryo Nitta. CAM-SAP2 organizes a γ-tubulin-independent microtubule nucleation centre through phase separation. eLife, 11:1–28, 2022. ISSN 2050084X. doi: 10.7554/eLife.77365.

25. Hao Jiang, Shusheng Wang, Yuejia Huang, Xiaonan He, Honggang Cui, Xueliang Zhu, and Yixian Zheng. Phase Transition of Spindle-Associated Protein Regulate Spindle Apparatus Assembly. Cell, 163(1):108–122, 2015. ISSN 10974172. doi: 10.1016/j.cell.2015.08.010.

26. Wanqing Lyu, Daisy Duan, Kuanlin Wu, Chunxiang Wu, Yong Xiong, and Anthony J Koleske. Ab12 mediates microtubule nucleation and repair via tubulin co-condensation. bioRxiv, page 2022.06.21.496973, 2022.

27. Yueh-Fu O Wu, Annamarie T Bryant, Nora T Nelson, Alexander G Madey, Gail F Fernandes, and Holly V Goodson. Overexpression of the microtubule-binding protein clip-170 induces a+tip network superstructure consistent with a biomolecular condensate. PloS one, 16(12): e0260401, 2021.

28. Julie Miesch, Robert T Wimbish, Marie-Claire Velluz, and Charlotte Aumeier. Phase separation of +tip-networks regulates microtubule dynamics. BioRxiv, pages 2021–09, 2022.

29. Kenneth J. Rosenberg, Jennifer L. Ross, H. Eric Feinstein, Stuart C. Feinstein, and Jacob Israelachvili. Complementary dimerization of microtubule-associated tau protein: Implications for microtubule bundling and tau-mediated pathogenesis. Proceedings of the National Academy of Sciences of the United States of America, 105(21):7445–7450, 2008. ISSN 00278424. doi: 10.1073/pnas.0802036105.

30. Xuemei Zhang, Yanxian Lin, Neil A. Eschmann, Hongjun Zhou, Jennifer Rauch, Israel Hernandez, Elmer Guzman, Kenneth S. Kosik, and Songi Han. RNA Stores Tau Reversibly in Complex Coacervates. PLoS Biology, 15(7):1–28, 2017. ISSN 1544-9173. doi: 10.1101/111245.

31. Nicholas M. Kanaan, Chelsey Hamel, Tessa Grabinski, and Benjamin Combs. Liquid-liquid phase separation induces pathogenic tau conformations in vitro. Nature Communications, 11(1), 2020. ISSN 20411723. doi: 10.1038/s41467-020-16580-3.

32. Tonglin Mao, Lifeng Jin, Hua Li, Bo Liu, and Ming Yuan. Two microtubule-associated proteins of the Arabidopsis MAP65 family function differently on microtubules. Plant Physiology, 138(2):654–662, 2005. ISSN 00320889. doi: 10.1104/pp.104.052456.

33. Graham M. Burkart and Ram Dixit. Microtubule bundling by MAP65-1 protects against severing by inhibiting the binding of katanin. Molecular Biology of the Cell, 30(13):1587–1597, 2019. ISSN 19394586. doi: 10.1091/mbc.E18-12-0776.

34. Bianca Edozie, Sumon Sahu, Miranda Pitta, Anthony Englert, Carline Fermino Do Rosario, and Jennifer L. Ross. Self-organization of spindle-like microtubule structures. Soft Matter, 15(24):4797–4807, 2019. ISSN 17446848. doi: 10.1039/c8sm01835a.

35. Sumon Sahu, Lena Herbst, Ryan Quinn, and Jennifer L. Ross. Crowder and surface effects on self-organization of microtubules. Physical Review E, 103(6):1–20, 2021. ISSN 24700053. doi: 10.1103/PhysRevE.103.062408.

36. Simon Alberti, Amy Gladfelter, and Tanja Mittag. Considerations and Challenges in Studying Liquid-Liquid Phase Separation and Biomolecular Condensates. Cell, 176(3):419–434, 2019. ISSN 10974172. doi: 10.1016/j.cell.2018.12.035.

37. Avinash Patel, Hyun O. Lee, Louise Jawerth, Shovamayee Maharana, Marcus Jahnel, Marco Y. Hein, Stoyno Stoynov, Julia Mahamid, Shambaditya Saha, Titus M. Franzmann, Andrej Pozniakovski, Ina Poser, Nicola Maghelli, Loic A. Royer, Martin Weigert, Eugene W. Myers, Stephan Grill, David Drechsel, Anthony A. Hyman, and Simon Alberti. A Liquid-to-Solid Phase Transition of the ALS Protein FUS Accelerated by Disease Mutation. Cell, 162 (5):1066–1077, 2015. ISSN 10974172. doi: 10.1016/j.cell.2015.07.047.

38. Tina W. Han, Masato Kato, Shanhai Xie, Leeju C. Wu, Hamid Mirzaei, Jimin Pei, Min Chen, Yang Xie, Jeffrey Allen, Guanghua Xiao, and Steven L. McKnight. Cell-free formation of RNA granules: Bound RNAs identify features and components of cellular assemblies. Cell, 149(4):768–779, 2012. ISSN 10974172. doi: 10.1016/j.cell.2012.04.016.

39. Masato Kato, Tina W. Han, Shanhai Xie, Kevin Shi, Xinlin Du, Leeju C. Wu, Hamid Mirzaei, Elizabeth J. Goldsmith, Jamie Longgood, Jimin Pei, Nick V. Grishin, Douglas E. Frantz, Jay W. Schneider, She Chen, Lin Li, Michael R. Sawaya, David Eisenberg, Robert Tycko, and Steven L. McKnight. Cell-free formation of RNA granules: Low complexity sequence domains form dynamic fibers within hydrogels. Cell, 149(4):753–767, 2012. ISSN 10974172. doi: 10.1016/j.cell.2012.04.017.

40. Amandine Molliex, Jamshid Temirov, Jihun Lee, Maura Coughlin, Anderson P. Kanagaraj, Hong Joo Kim, Tanja Mittag, and J. Paul Taylor. Phase Separation by Low Complexity Domains Promotes Stress Granule Assembly and Drives Pathological Fibrillization. Cell, 163(1):123–133, 2015. ISSN 10974172. doi: 10.1016/j.cell.2015.09.015.

41. Ilmin Kwon, Masato Kato, Siheng Xiang, Leeju Wu, Pano Theodoropoulos, Hamid Mirzaei, Tina Han, Shanhai Xie, Jeffry L. Corden, and seven L. McKnight. XPhosphorylation-regulated binding of RNA polymerase II to fibrous polymers of low-complexity domains. Cell, 155(5):1049, 2013. ISSN 10974172. doi: 10.1016/j.cell.2013.10.033.

42. Siheng Xiang, Masato Kato, Leeju C. Wu, Yi Lin, Ming Ding, Yajie Zhang, Yonghao Yu, and Steven L. McKnight. The LC Domain of hnRNPA2 Adopts Similar Conformations in Hydrogel Polymers, Liquid-like Droplets, and Nuclei. Cell, 163(4):829–839, 2015. ISSN 10974172. doi: 10.1016/j.cell.2015.10.040.

43. Marina Feric, Nilesh Vaidya, Tyler S. Harmon, Diana M. Mitrea, Lian Zhu, Tiffany M. Richardson, Richard W. Kriwacki, Rohit V. Pappu, and Clifford P. Brangwynne. Coexisting Liquid Phases Underlie Nucleolar Subcompartments. Cell, 165(7):1686–1697, 2016. ISSN 10974172. doi: 10.1016/j.cell.2016.04.047.

44. Tetsuro Murakami, Seema Qamar, Julie Qiaojin Lin, Gabriele S. Kaminski Schierle, Eric Rees, Akinori Miyashita, Ana R. Costa, Roger B. Dodd, Fiona T.S. Chan, Claire H. Michel, Deborah Kronenberg-Versteeg, Yi Li, Seung Pil Yang, Yosuke Wakutani, William Meadows, Rodylyn Rose Ferry, Liang Dong, Gian Gaetano Tartaglia, Giorgio Favrin, Wen Lang Lin, Dennis W. Dickson, Mei Zhen, David Ron, Gerold Schmitt-Ulms, Paul E. Fraser, Neil A. Shneider, Christine Holt, Michele Vendruscolo, Clemens F. Kaminski, and Peter St George-Hyslop. ALS/FTD Mutation-Induced Phase Transition of FUS Liquid Droplets and Reversible Hydrogels into Irreversible Hydrogels Impairs RNP Granule Function. Neuron, 88(4):678–690, 2015. ISSN 10974199. doi: 10.1016/j.neuron.2015.10.030.

45. Hong Joo Kim, Nam Chul Kim, Yong Dong Wang, Emily A. Scarborough, Jennifer Moore, Zamia Diaz, Kyle S. MacLea, Brian Freibaum, Songqing Li, Amandine Molliex, Anderson P. Kanagaraj, Robert Carter, Kevin B. Boylan, Aleksandra M. Wojtas, Rosa Rademakers, Jack L. Pinkus, Steven A. Greenberg, John Q. Trojanowski, Bryan J. Traynor, Bradley N. Smith, Simon Topp, Athina Soragia Gkazi, Jack Miller, Christopher E. Shaw, Michael Kottlors, Janbernd Kirschner, Alan Pestronk, Yun R. Li, Alice Flynn Ford, Aaron D. Gitler, Michael Benatar, Oliver D. King, Virginia E. Kimonis, Eric D. Ross, Conrad C. Weihl, James Shorter, and J. Paul Taylor. Mutations in prion-like domains in hnRNPA2B1 and hnRNPA1 cause multisystem proteinopathy and ALS. Nature, 495(7442):467–473, 2013. ISSN 00280836. doi: 10.1038/nature11922.

46. Louise Jawerth, Elisabeth Fischer-Friedrich, Suropriya Saha, Jie Wang, Titus Franzmann, Xiaojie Zhang, Jenny Sachweh, Martine Ruer, Mahdiye Ijavi, Shambaditya Saha, Julia Mahamid, Anthony A. Hyman, and Frank Jülicher. Protein condensates as aging Maxwell fluids. Science, 370(6522):1317–1323, 2020. ISSN 10959203. doi: 10.1126/science.aaw4951.

47. Timothy J. Nott, Evangelia Petsalaki, Patrick Farber, Dylan Jervis, Eden Fussner, Anne Plochowietz, Timothy D. Craggs, David P. Bazett-Jones, Tony Pawson, Julie D. Forman-Kay, and Andrew J. Baldwin. Phase Transition of a Disordered Nuage Protein Generates Environmentally Responsive Membraneless Organelles. Molecular Cell, 57(5):936–947, 2015. ISSN 10974164. doi: 10.1016/j.molcel.2015.01.013.

48. Shana Elbaum-Garfinkle, Younghoon Kim, Krzysztof Szczepaniak, Carlos Chih Hsiung Chen, Christian R. Eckmann, Sua Myong, and Clifford P. Brangwynne. The disordered P granule protein LAF-1 drives phase separation into droplets with tunable viscosity and dynamics. Proceedings of the National Academy of Sciences of the United States of America, 112(23):7189–7194, 2015. ISSN 10916490. doi: 10.1073/pnas.1504822112.

49. Minchul Kang, Charles A. Day, Anne K. Kenworthy, and Emmanuele DiBenedetto. Simplified equation to extract diffusion coefficients from confocal FRAP data. Traffic, 13(12):1589–1600, 2012. ISSN 13989219. doi: 10.1111/tra.12008.

50. Jérémie Gaillard, Emmanuelle Neumann, Daniel Van Damme, Virginie Stoppin-Mellet, Christine Ebel, Elodie Barbier, Danny Geelen, and Marylin Vantard. Two Microtubule-associated Proteins of Arabidopsis MAP65s Promote Antiparallel Microtubule Bundling. Molecular Biology of the Cell, 19(10):4534–4544, 2008. doi: 10.1091/mbc.e08-04-0341.

51. Jeffrey B. Woodruff, Beatriz Ferreira Gomes, Per O. Widlund, Julia Mahamid, Alf Honig-mann, and Anthony A. Hyman. The Centrosome Is a Selective Condensate that Nucleates Microtubules by Concentrating Tubulin. Cell, 169(6):1066–1077, 2017. ISSN 10974172. doi: 10.1016/j.cell.2017.05.028.

52. Haoge Li, Baojuan Sun, Michiko Sasabe, Xingguang Deng, Yasunori Machida, Honghui Lin, Y. R. Julie Lee, and Bo Liu. Arabidopsis MAP65-4 plays a role in phragmoplast microtubule organization and marks the cortical cell division site. New Phytologist, 215(1):187–201, 2017. ISSN 14698137. doi: 10.1111/nph.14532.

53. Andrei P Smertenko, Despina Kaloriti, Hsin-Yu Chang, Jindriska Fiserova, Zdenek Opatrny, and Patrick J Hussey. The c-terminal variable region specifies the dynamic properties of arabidopsis microtubule-associated protein map65 isotypes. The Plant Cell, 20(12):3346–3358, 2008.

54. Jessica R. Lucas, Stephanie Courtney, Mathew Hassfurder, Sonia Dhingra, Adam Bryant, and Sidney L. Shaw. Microtubule-associated proteins MAP65-1 and MAP65-2 positively regulate axial cell growth in etiolated Arabidopsis hypocotyls. Plant Cell, 23(5):1889–1903, 2011. ISSN 10404651. doi: 10.1105/tpc.111.084970.

55. Kimberly L. Weirich, Kinjal Dasbiswas, Thomas A. Witten, Suriyanarayanan Vaikuntanathan, and Margaret L. Gardel. Self-organizing motors divide active liquid droplets. Proceedings of the National Academy of Sciences, 116(23):11125–11130, 2019. doi: 10.1073/pnas.1814854116.

56. Kimberly L. Weirich, Shiladitya Banerjee, Kinjal Dasbiswas, Thomas A. Witten, Suriyanarayanan Vaikuntanathan, and Margaret L. Gardel. Liquid behavior of cross-linked actin bundles. Proceedings of the National Academy of Sciences of the United States of America, 114(9):2131–2136, 2017. ISSN 10916490. doi: 10.1073/pnas.1616133114.

57. Danielle R. Scheff, Kimberly L. Weirich, Kinjal Dasbiswas, Avinash Patel, Suriyanarayanan Vaikuntanathan, and Margaret L. Gardel. Tuning shape and internal structure of protein droplets via biopolymer filaments. Soft Matter, 16(24):5659–5668, 2020. ISSN 1744-683X. doi: 10.1039/c9sm02462j.

58. Kristin Graham, Aravind Chandrasekaran, Liping Wang, Aly Ladak, Eileen M Lafer, Padmini Rangamani, and Jeanne C Stachowiak. Liquid-like VASP condensates drive actin polymerization and dynamic bundling. bioRxiv, page 2022.05.09.491236, 1 2022. doi: 10.1101/2022.05.09.491236.

59. Lindsay B. Case, Xu Zhang, Jonathon A. Ditlev, and Michael K. Rosen. Stoichiometry controls activity of phase-separated clusters of actin signaling proteins. Science, 363(6431): 1093–1097, 2019. ISSN 10959203. doi: 10.1126/science.aau6313.

60. Sen Yang, Chunxia Liu, Yuting Guo, Guoqing Li, Dong Li, Xiumin Yan, and Xueliang Zhu. Self-construction of actin networks through phase separation-induced abLIM1 condensates. Proceedings of the National Academy of Sciences of the United States of America, 119(29), 2022. ISSN 10916490. doi: 10.1073/pnas.2122420119.

61. Ying Xie, Jialin Sun, Xiao Han, Alma Turšić-Wunder, Joel D.W. Toh, Wanjin Hong, Yong Gui Gao, and Yansong Miao. Polarisome scaffolder Spa2-mediated macromolecular condensation of Aip5 for actin polymerization. Nature Communications, 10(1), 2019. ISSN 20411723. doi: 10.1038/s41467-019-13125-1.

62. Tina Wiegand and Anthony A. Hyman. Drops and fibers -How biomolecular condensates and cytoskeletal filaments influence each other. Emerging Topics in Life Sciences, 4(3): 247–261, 2020. ISSN 23978562. doi: 10.1042/ETLS20190174.

