## Supplemental Data for "Spatially controlled microtubule nucleation and organization from crosslinker MAP65 condensates"

### Materials and Methods

All chemicals were purchased from Sigma unless otherwise noted.

#### Protein purification.

***GFP-MAP65 and MAP65 purification.*** The microtubule crosslinker, MAP65-1 (MAP65) plasmid was a gift from Ram Dixit (Washington University, St. Louis). The full protein purification protocol is detailed in (1, 2). Briefly, protein is expressed using *Escherichia coli* BL21(DE3) cells, grown to an $OD_{600}$ of 1, lysed with sonication, and clarified via centrifugation. MAP65 protein is recovered from the lysate through affinity between the 6× histidine tag and Ni-binding substrate (1). Purified protein was buffer exchanged to PEM80, checked on an SDS-PAGE gel to measure purification and concentration, aliquoted, and stored at -80 $^oC$.

***GFP-PRC1 and PRC1 purification.*** The WT full length PRC1 plasmid was a gift from Radhika Subramanian (Massachusetts General Hospital). The detailed purification protocol can be found in (3). Briefly, protein is expressed using *Escherichia coli* BL21(DE3) Rosetta cells, grown to an $OD_{600}$ of 0.6-0.7 in LB media before inducing with IPTG. After the cells were grown at $18^oC$, they were harvested via centrifugation and resuspended in lysis buffer with TCEP, Pierce Protesase inhibitor, PMSF, and Benz HCl. The cells were further sonicated and centrifuged. The PRC1 protein was recovered from the lysate through affinity between the 6× histidine tag and Ni-NTA substrate. Purified protein was dialyzed, passed through size exclusion column, and concentrated using Amicon Ultra centrifugal filter. The protein was then aliquoted, and stored at -80 $^oC$ with 30% sucrose.

**Tubulin preparation.** Unlabeled and fluorescently labeled (rhodamine) lyophilized tubulin from porcine brain is purchased from Cytoskeleton. Tubulins are resuspended to 5 mg/ml in PEM-80 buffer (80 mM PIPES, pH 6.8, 1 mM $MgCl_2$, 1 mM EGTA). Fluorescent and unlabeled tubulin are combined to a final labeling ratio of $\sim$ 10% fluorescently labeled tubulin. Tubulin is aliquoted, drop-frozen, and stored in -80 $^oC$ for later use. Aliquots are thawed on ice prior to use.

### LLPS assays

**Silanized coverslips.** Coverslips (Corning, 22 mm × 30 mm × 0.17 mm) are treated using a block co-polymer brush of Pluronic-F127, as previously described to inhibit protein binding to the surface (4, 5). Coverslips are cleaned with ethanol, acetone, potassium hydroxide (KOH) from Sigma, and treated with 2% (w/v) dimethyldichlorosilane solution (Cytiva) to make the surface hydrophobic. The full silanization protocol can be found in (6).

**Sample preparation and flow chamber.** GFP-MAP65 and unlabeled MAP65 were mixed together to make 10% labeled MAP65 in PEM80 (80 mM PIPES, 1 mM EGTA, 2 mM $MgCl_2$) in PEM80. For PRC1 experiments, GFP-PRC1 and unlabeled PRC1 were mixed together to make 10% labeled PRC1 in PRC1 storage buffer (50 mM NaP pH 7.0, 150 mM NaCl). The sample was kept in room temperature for 20-25 min. In the meantime, a 10 $\mu$l flow chamber was made using glass slide (Fisher), silanized coverslip, and double sided tape (3M). A solution of 5% F127 in water was added to the chamber and incubated for 7-10 min. Once the sample incubation is over, the sample was flowed into the chamber using capillary flow by filter paper and was sealed with epoxy on both ends. The chamber was kept upside down for 5-15 min in order to let the condensates settle by gravity. Initial videos were taken right after that to capture droplet coalescence events. The images for analysis were taken approximately 45-60 minutes after the mix. We also performed experiments with samples where dithiothreitol (DTT) is present and do not find significant difference in droplet formation. However, we have observed that MAP65 condensates age faster becoming gel-like when an oxygen scavenging system consisting of glucose oxidase and catalase, DTT, and glucose were present. In order to have fluid-like properties longer, we eliminated the oxygen scavenging system from our experiments.

For aster-formation experiments, a cloning glass cylinder (Fisher: 0955221) of dimension 8 mm × 8 mm was used. The cylinder was cleaned with water, ethanol, and water, dried and was attached to a silanized coverslip (Corning, 22 mm × 30 mm × 0.17 mm) with epoxy. The chamber was incubated with 40 $\mu l$ F127 for 10 minutes. After that F127 was pipeted out and tubulin sample (with GMPcPP or no GMPcPP) from ice was added to the chamber. Then, already formed MAP65 condensate solution (at room temperature) was added to the chamber. Finally, 10 $\mu l$ mineral oil was added to the chamber to stop evaporation. After this point sample was at room temperature while asters formed. The imaging was performed starting from after an hour of sample addition.

### Confocal imaging and photobleaching experimental procedures

**Confocal imaging.** Imaging of labeled condensates is performed using spinning disc microscopy (Yokogawa CSU-W1 50um Pinhole) on an inverted Nikon Ti-E microscope with Perfect Focus and 100x oil immersion objective (1.49 NA) imaged onto a Andor Zyla CMOS camera. Image acquisition is performed with 488 laser and occasionally with 561 laser. The image scale is 65 nm per pixel. Images are displayed and recorded using the Nikon Elements software. Time series data was saved as .nd2 files with metadata and analyzed using ImageJ/FIJI and MATLAB.

**Photobleaching.** Photobleaching experiments are performed by imaging with the 488 nm and 561 nm lasers and photobleaching with a 405 nm laser using an Optomicroscan Laser FRAP module attached to the Nikon Ti-E microscope. The dot photobleaching data was taken as follows: 1) Acquisition: 16 seconds with 2 seconds interval, 2) Photobleaching: 20 milliseconds, 3) Acquisition: 120 seconds with 2 seconds interval. Four point ROIs were selected in Nikon elements software to photobleach the designated regions in the condensates in the same timeseries. Time series data was saved as .nd2 files with metadata and analyzed using ImageJ/FIJI. The whole droplet photobleaching data was taken as follows: 1) Acquisition: 16 seconds with 2 seconds interval, 2) Photobleaching: 1.73 seconds, 3) Acquisition: 300 seconds with 2 seconds interval. Four square ROIs were selected in Nikon elements software to photobleach the designated regions.

### Image analysis

For image analysis we have used ImageJ/FIJI and custom MATLAB scripts. In MATLAB scripts, all raw images were pre-processed in the following manner. First, we applied $4 * 4$ median filter to get rid of the noise present in the image. Then, the image was binarized into black and white. Further, to get rid of boundary objects in-built function ‘imclearborder’ was used followed by ‘bwareafilt’ to filter out very small objects which are noise. To extract features, we have called in-built ‘regionprops’ function in MATLAB.

**Intensity quantification: line scans to measure partition coefficient.** To calculate intensity distribution over a line, we used line tool and plot profile in ImageJ/FIJI. The images selected was a confocal slice through the middle of the droplet. The average intensity within the bright droplet region was calculated from the region of the line on the droplet. The background intensity was determined from the line scan of the edges outside the droplet. The intensity data inside the droplet was divided by the intensity outside the droplet to determine the partition co-efficient for a line scan.

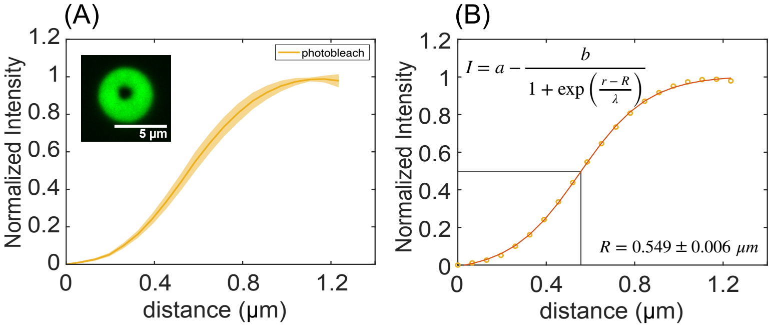


**Fig. S1.** Measurement of photobleached radius: (A) Radially averaged normalized intensity profile of a photobleached spot is shown (n=16). The solid line (yellow) indicates the mean, and the shaded area (light yellow) indicates the standard deviation. Inset - A representative droplet with a point photobleached spot is shown. (B) The radially averaged normalized mean profile is fitted with the given sigmoid equation. The radius of the photobleached spot, $R \sim 0.55\ \mu m$, is obtained from the fit. R signifies the point where the normalized intensity is 0.5.

**Intensity quantification: whole droplet intensity and partition co-efficient.** The mean intensity of the background was calculated manually by selecting an ROI in ImgeJ/FIJI that was not overlapping any droplets. The mean intensity of each droplet was calculated using ‘regionprops’ function in MATLAB after pre-processing. Then these mean intensities were averaged to get an interior average mean intensity and background average mean intensity (Fig. S2(D) inset left and right). To calculate the partition co-efficient, the ratio of interior and background average mean intensity was calculated. SEM was calculated as error bars accordingly.

**Fluorescence recovery after photobleaching analysis.** MATLAB was used to analyze the FRAP data. For point photobleaching, we calculated the mean intensity of the photobleached region within the droplet by selecting a circular ROI in Nikon elements software and stored as a raw intensity data. The radius of the ROI was determined by taking a radial average from the center of the photobleached spot and fit in MATLAB (n=16) using $I = a - \frac{b}{1+exp(\frac{r-R}{\lambda})}$. Here, fitting parameters a and b are typically 1, R is the radius at which $I = 0.5$, and $\lambda$ is the length scale (Fig. S1). To correct for global photobleaching, the mean intensity values from another same-radius ROI on a non-photobleached droplet were also recorded (7). For, whole droplet photobleaching or ring shaped photobleaching, radially averaged intensity was calculated in MATLAB and further averaging was performed to the condensate boundary. For aster photobleaching, mean intensity was calculated in FIJI using an ROI on the aster. The corrected raw curves were normalized and averaged for each data set. We fit the normalized photo recovery mean curve with the given equation (8, 9),

$$I(t) = \frac{a + \frac{bt}{\tau_{1/2}}}{1 + \frac{t}{\tau_{1/2}}} \qquad [1]$$

where I(t) is the intensity value at time t, $\tau_{1/2}$ is the recovery constant, and $a$ and $b$ are the fitting parameters.

**Volume quantification.** To check if the volume of the droplet is conserved before and after coalescence, we performed measurements manually in ImageJ/FIJI. The diameter, $D$, of each droplet was measured five times for each droplet through the center of the droplet using the line tool and the volume was approximated using $v = 4/3\pi r^3$ where $r = D/2$. Then mean and standard deviation was calculated for droplet volumes $V_1$ and $V_2$ before coalescence, and $V_3$ after coalescence. Images from both bright-field imaging and confocal microscopy were used to calculate volume. The images used were taken at the mid-point of the droplet to accurately reflect the largest volume. The volume of droplets before and after coalescence were plotted and a linear fit was used to fit the data in the main text.

**Aspect ratio quantification.** We calculated the aspect ratio of two droplets as they coalesced over time using confocal image stacks processed using a custom MATLAB script. After pre-processing, an ellipsoid was fit to the object of interest to get major axis length l, minor axis length w, and aspect ratio $l/w$. We have chosen nearly identical sized droplets for this analysis. The measured aspect ratio over time is fit to the $1 + a\ \exp(-t/\tau_e)$ equation to get the relaxation time $\tau_e$.

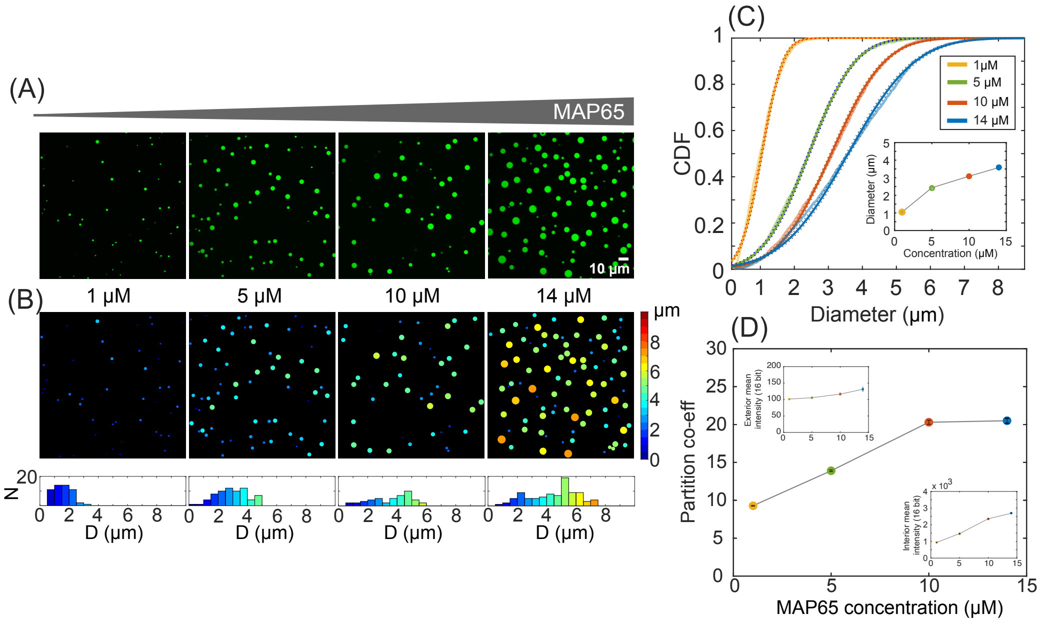


**Fig. S2.** (A) Representative raw images are shown for an increasing MAP65 concentration, [MAP65]= $1\ \mu M$, $5\ \mu M$, $10\ \mu M$, and $14\ \mu M$ from left to right. Larger number densities with higher droplet diameters are observed from left to right. (B) Droplets sorted according to their diameter and color-coded to visualize the respective images. Bottom - The histograms represent diameter distribution for those representative images, which shifts from narrow mono-disperse distribution to wide distribution as we increase the MAP65 concentration. (C) Cumulative distribution function of condensate diameters (1) $1\ \mu M$ (yellow, n=1564), (2) $5\ \mu M$ (green, n=2716), (3) $10\ \mu M$ (red, n=2632), (4) $14\ \mu M$ (blue, n=2466). The lighter colors represent the data, the darker colors of the same shade represent the normal distribution fit to the data, and the dots represent the $95\ \%$ confidence interval to the fit. (c-inset) The mean diameter calculated from the fitted distribution with standard error of the mean (SEM) is plotted against MAP65 concentration. The fitted diameter values $(mean \pm SEM)$ are (1) D = $1.04 \pm 0.01\ \mu m$ for $1\ \mu M$, (2) D = $2.43 \pm 0.02\ \mu m$ for $5\ \mu M$, (3) D = $3.08 \pm 0.02\ \mu m$ for $10\ \mu M$, (4) D = $3.58 \pm 0.03\ \mu m$ for $14\ \mu M$. (D) Partition co-efficient value $p = I_{in}/I_{out}$ (mean $\pm$ SEM)increases and finally saturates with increased MAP65 concentration, (1) $1\ \mu M$ (yellow, n=1564), (2) $5\ \mu M$ (green, n=2716), (3) $10\ \mu M$ (red, n=2632), (4) $14\ \mu M$ (blue, n=2466). For each concentration, average intensity of background was measured (left-inset) and average intensity of each droplet was measured (right-inset).

### Supplemental experiments

**MAP65 concentration effect on condensates.** To check the effect of MAP65 concentration on droplet formation, we performed experiments with four different concentrations, $1\ \mu M$, $5\ \mu M$, $10\ \mu M$, and $14\ \mu M$. We observed that the condensate number and size/diameter grew as the MAP65 concentration increased. In addition, the surface fraction, the area covered by droplets to the area of the viewing window, increases (Fig. S2A). For each concentration, we analyzed at least n=1500 droplets from at least 28 raw images to quantify the condensate diameter values. These images were taken under the same conditions and within a similar time window as described. MATLAB codes were used to automatically identify droplets, measure their size, and color the droplets based on size (Fig. S2B).

We created the cumulative probability distributions (CDF) of the droplet diameters to compare the different MAP65 concentrations. We find that the distributions of the diameters display a normal distributions (Fig. S2B, C). We fit the CDF with Gaussian functions to find the mean, $\mu$, and standard deviation, $\sigma$, of the best fits. The mean of the droplet diameters shows an upward trend as the MAP65 concentration increases (Fig. S2C-inset). This result implies that bigger droplets are formed with more MAP65. It is likely that these larger droplets are the result of coalescing many smaller droplets. When the MAP65 concentration is high, we expect that a high density of smaller condensates are nucleated. Due to their small state, these droplets diffuse faster. Due to their liquid-like state, the droplets can fuse into larger droplets when they meet. At higher number densities, the chance of encountering another droplet is higher in the initial times after flowing the sample when droplets are mobile. This causes larger condensates at higher concentrations.

For each concentration of MAP65, we also measured the partition coefficient using the methods described above (Fig. S2D). We find that as the concentration of MAP65 increases, both the intensity of the fluorescence outside (Fig. S2D, inset top left) and inside the condensed droplets increases (Fig. S2D, inset bottom right). The ratio of these intensities is the partition coefficient, which increases as well since the intensity inside the droplets is higher at higher MAP65 concentrations (Fig. S2(D). Thus, more MAP65 is in the droplets with higher MAP65 concentrations.

**PRC1 condensate formation in vitro.** The full-length PRC1 domains with C-terminal intrinsically disordered region (IDR) are shown in Fig. S3A (3). The two different PONDR algorithms (VSL3 and VL3) predict the PRC1 disordered region. When compared to MAP65, PRC1 also has locally concentrated positive charge regions in the C-terminal tail which is predicted by CIDER net charge per residue (NCPR) distribution (Fig. S3A).

Indeed, purified full length WT PRC1 phase separates at room temperature $22 \pm 1\ ^{o}C$ in phosphate buffer (Fig. S3B). We confirmed both unlabeled PRC1 and 10% GFP-PRC1 with unlabled PRC1 phase separate by bright field microscopy and confocal microscopy, respectively (Fig. S3B). We confirmed that PRC1 phase separates in PEM-80, as well.

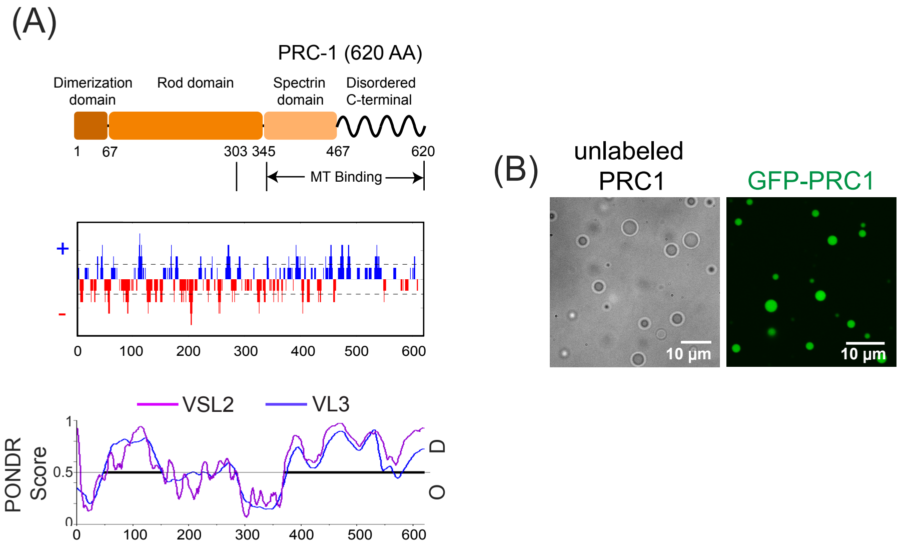


**Fig. S3.** (A) Full length WT PRC1 regions - dimerization domain(dark orange), rod domain (orange), spectrin domain (lighter orange), and LCD are shown by boxes and a wavy line, respectively. The predicted net charge per residue (NCPR) distribution from CIDER shows where blue and red represent positive and negative charges, respectively. PONDR algorithms, VSL2 and VL3, predict disorder probability for PRC1. (B) Left - Bright-field image shows unlabeled full-length PRC1 forms condensates. Right - 10% labeled GFP-PRC1 condensates in confocal 488 channel.

**Salt and temperature effects on MAP65 and PRC1 condensates.** We have tested the effect of additional monovalent salt and temperature conditions on the MAP65 and PRC1 droplets. Addition of KCl and NaCl for MAP65 and PRC1 condensates, respectively inhibited the condensate formation (Fig. S4Ai-Bi). Both MAP65 and PRC1 has localized positively charged disordered C-terminal tail. This result may suggest that MAP65-MAP65 or PRC1-PRC1 interactions are charged interactions and happen through the disordered tail.

We have also tested the temperature effects on MAP65 and PRC1 condensates. At low temperature, 4 $^{o}C$, both formed small condensates. At human body temperature, 37 $^{o}C$, while MAP65 condensates were aged and made deformed structures, PRC1 was round liquid-like and were able to coalesce (Fig. S4Aii-Bii).

**Whole droplet photobleaching and estimation of diffusion co-efficient.** We performed fluorescence recovery after photobleaching to measure the diffusion of MAP65 within the condensate and the exchange at the boundary of the condensate. In order to understand exchange at the condensate boundary, we photobleached entire droplets and watched the recovery of the GFP-labeled MAP65 from outside the droplet, as diagrammed in the cartoon (Fig. S5(A)). To measure the intensity in the droplet, we quantified and plotted the radial average as a function of distance from the center of the droplet (Fig. S5(A)).

Whole droplet fluorescence recovery was measured as a function of droplet age at time points of 1.5 h, 2.5 h, 3.5 h, and 4.5 h (Fig. S5(B)). At early times, the droplet intensity recovers within 300 s, although never to the initial level. As the droplets age, the mobile fraction molecules decrease due to the increased viscosity inside the droplets. Interesting, it is visible that the new GFP-MAP65 added to the droplets is coming from the exterior, as there is a ring of bright, labeled MAP65 molecules from

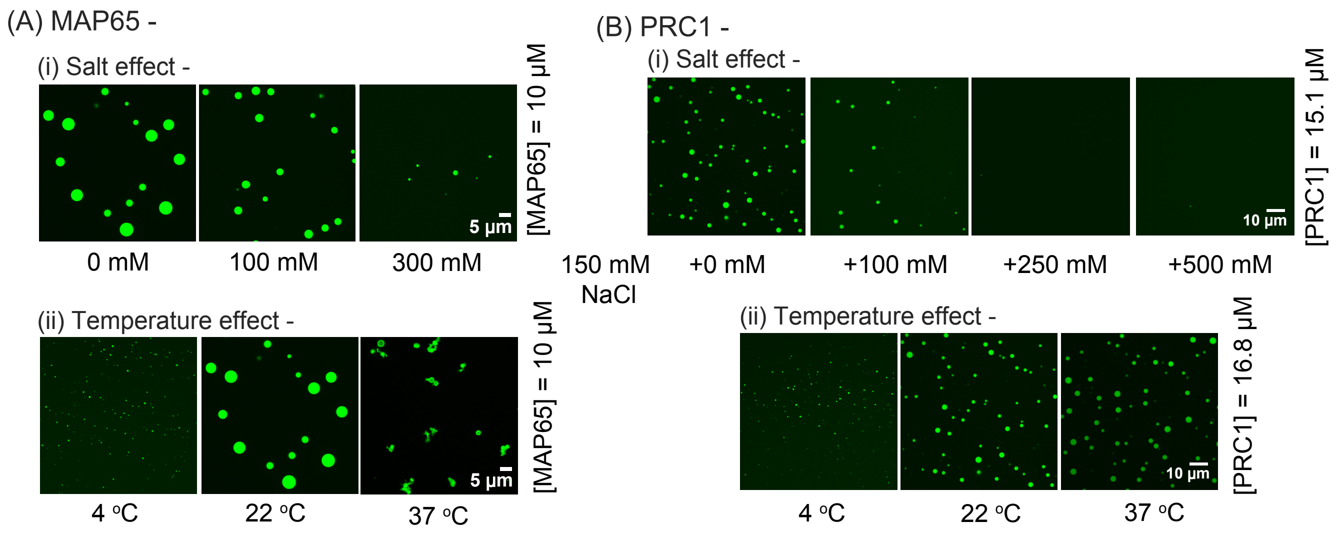


**Fig. S4.** (A)(i) Representative images from three configurations with KCl salt concentrations [KCl]= $0\ mM,\ 100\ mM$, and $300\ mM$ with [MAP65]= $10\ \mu M$ show inhibition of condensate formation as the KCl concentration is increased. Very few to almost no droplets are formed at the highest KCl concentration. The scale bar is $10\ \mu m$. (ii) Representative images from three different experiments with varied temperature T=$4\ ^oC,\ 22\ ^oC$ and $37\ ^oC$ are shown. While at $4\ ^oC$, small condensates are formed too, at $37\ ^oC$ already formed condensates quickly aged into the viscoelastic droplets that create a chain/network of jammed droplets. (B)(i) Representative images from four configurations with added NaCl salt concentrations [NaCl]= $0\ mM,\ 100\ mM, 250\ mM$, and $500\ mM$ with [PRC1]= $15.1\ \mu M$ show inhibition of condensate formation as the NaCl concentration is increased. Almost no droplets are formed at the highest NaCl concentration. The scale bar is $10\ \mu m$.(ii) Representative images from three different experiments with varied temperature T=$4\ ^oC,\ 22\ ^oC$ and $37\ ^oC$ are shown. At $37\ ^oC$ are still round shaped and liquid-like.

outside the droplet. We used the radially averaged intensity profiles over time, $I(r,t)$ to show the change in dynamics over time (Fig. S5B). From normalized $I(r,t)$, $I_n(r,t)$ we also noticed that the material exchange at the boundary is also decreased with T when measured at t = 10 sec and t = 290 sec after photobleaching for every data points (Fig. S5C i-ii). This may suggest that the availability of GFP-MAP65 is also depleted over time in the exterior as most of them are partitioned into other droplets as the exchange dynamics is slowed down. In another way, this reports the rate of molecule exchange between the droplet and the surrounding MAP65 pool.

We can use the normalized radial intensity scans over time, $I_n(r,t)$, combined with knowledge of the basics of diffusion to directly solve the differential equations governing the processes of MAP65 diffusion in the droplets. Previously, whole droplet photobleaching analysis combined with the solution from diffusion equation is reported for protein/synthetic system (10). We have adapted and modified the MATLAB code to perform the MAP65 photobleaching analysis.

$$\frac{\partial I_n(r,t)}{\partial t} = -\nabla \cdot j \qquad [2]$$

$$j = -D_{in} \nabla I_n(r,t) \qquad [3]$$

Here, $I_n(r,t)$ is the normalized intensity of the droplet as a function of radius, $r$, and time, $t$. The index, $n$ denotes that the intensity was normalized. The function $j$ is the flux of the GFP-MAP65 molecules which is proportional to the gradient of concentration of the molecules, i.e. $I_n(r,t)$. The parameter $D_{in}$ is the diffusion coefficient we are interested to determine. Since there is azimuthal symmetry in the interior intensity recovery profile, we radially averaged the intensity profile, reducing the dimension of the problem to one dimension. The equation [2] becomes 1D diffusion equation in radial co-ordinate,

$$\frac{\partial I_n(r,t)}{\partial t} = \frac{D_{in}}{r}\frac{\partial}{\partial r}\left(r\frac{\partial I_n(r,t)}{\partial r}\right) \qquad [4]$$

$$= D_{in}\left[\frac{\partial^2 I_n(r,t)}{\partial^2 r} + \frac{1}{r}\frac{\partial I_n(r,t)}{\partial r}\right] \qquad [5]$$

We used MATLAB's partial differential equation solver, 'pdepe' to solve the equation [5]. The intensity values at a certain distance ($r = R_-$) from the actual interface (see Fig. S6A) were selected as the dynamical Dirichlet boundary condition for the equation. This is done to avoid artifacts due to imaging, including the point spread function and background photobleaching due to the illumination. As an initial condition, the radial intensity profile at $t = 10$ sec is fed into the equation. This was done to avoid the noise artifacts present just after photobleaching. We select eight frames from the photorecovery image data and fit them with the solution from equation [5] while varying the sole parameter $D_{in}$ (Fig. S6B). For each value of $D_{in}$, we calculated the cost function $\phi$ (Fig. S6C) as a measure of the goodness of fit to the experimental values. After fitting the $\phi$ with a spline function and using another MATLAB function, 'fminsearch' the best $D_{in}$ is determined from the minima of the $\phi$ cost function.

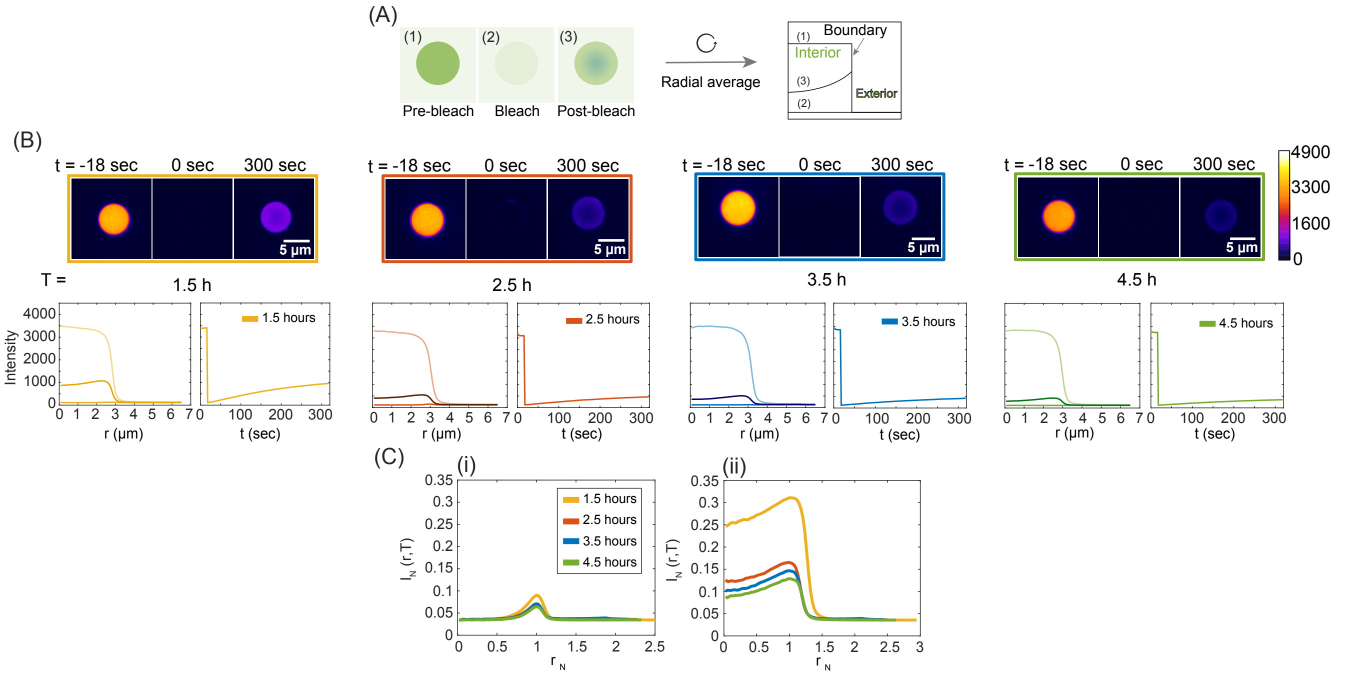


**Fig. S5.** Whole droplet photobleaching sample analysis - (A) Schematic diagram shows three time points of whole droplet photobleaching (1) pre-bleach, (2) bleach, and (3) post-bleach state for darker green condensate in lighter green background. After radial averaging, the cartoon plot shows intensity profiles of those three states. The ideal sharp boundary divide the interior and the exterior of the condensate. (B) Top - the representative images from four time points along the aging process, T = 1.5 hours (yellow box), 2.5 hours (red box), 3.5 hours (blue box), and 4.5 hours (green box) are shown in four columns. In each of the boxes the three images represent three photobleached states at time, t = -18 sec, 0 sec, and 300 sec. The completely photobleached state is denoted as time origin. The scale bars for all images are 5 $\mu m$. Bottom, left - at each time point T, a radially averaged raw intensity profile $I(r,t)$ is plotted at t=-18 sec, 0 sec, and 300 sec after photobleaching. Bottom, right - The mean intensity calculated using the radially averaged intensity plot from r=0 $\mu m$ to the droplet boundary is plotted with time t. (C) Normalized radially averaged intensity profile $I_N(r,t)$ is plotted against normalized r, $r_N$ to compare slightly different sized droplets at (i) t=10 sec and (ii) 290 sec at different time T = 1.5 hours (yellow), 2.5 hours (red), 3.5 hours (blue), and 4.5 hours (green).

We found that the best $D_{in}$ value fits the experimental normalized intensity profile $I_{N,exp}(r,t)$ to the solution of [5] well and also reconstructs the intensity at the $r = 0$ $\mu m$ or the left boundary (Fig. S6D-E).

**Table S1. MAP65 condensate aging data from whole condensate photobleaching**

| Time (h) | $D_{in} \times 10^{-2}$ $\mu m^2/s$ (mean $\pm$ std) |
|---|---|
| 1.5 | 1.18± 0.06 |
| 2.5 | 0.935± 0.005 |
| 3.5 | 0.84± 0.09 |
| 4.5 | 0.79± 0.07 |

When we compared the whole condensate photobleaching data to the dot photobleaching data described in the main text Table 1, the results are in good agreement within 2-2.5 hours after mixing while the droplets are in liquid-like state and show quicker dynamics. In later stages as the droplet property changes the results start to deviate where one overestimates the other. Although, overall, the aging dynamics is well captured in this whole condensate photobleaching analysis too Table S1.

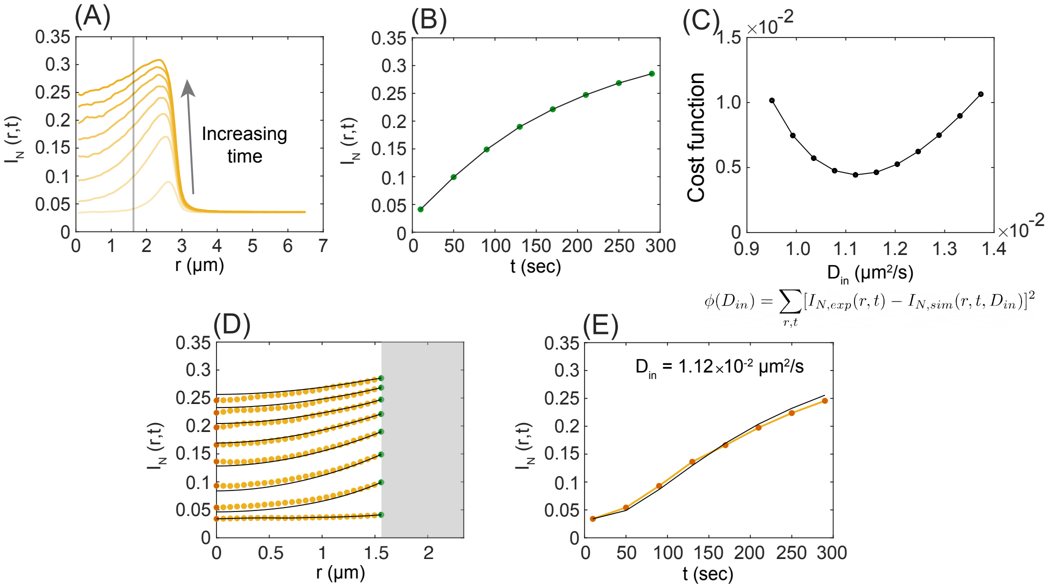


**Fig. S6.** Representative photobleaching analysis of a condensate at T=1.5 hours. (A) The radially averaged normalized intensity profile $I_N(r,t)$ is shown as a function of radial distance, r with increasing time, t. The vertical line represents the boundary which is used for further calculations. (B) The intensity values (green markers) at the vertical boundary $I_N(r = R_-, t)$ is used as dynamical boundary conditions in numerically solving equation [5], hence curves from experiment and fit superimpose on each other. (C) The diffusion co-efficient $D_in$ values are used as a parameter to fit the experimental intensity profile $I_{N,exp}(r,t)$ with the solution from equation [5], $I_{N,sim}(r,t)$. The parabola shaped local cost function, $\phi$ is plotted against trial $D_in$ values. (D) The solution (lines) at $D_in$ value resulted from $\phi$ minima are plotted with experimental values (dots). The left and right boundaries are indicated in red and green dots, respectively. (E) The experimental intensity values at the radial center (red markers, left boundary) with the solution from equation [5] with $D_in$ = 1.12 $\times 10^{-2}$ $\mu m^2/s$ is plotted against time t.

**Condensate surface tension estimate.** The complete coalescence of two MAP65 condensates can only occur if the droplets are liquid enough to fuse; viscoelastic droplets may bind and stick, but not have enough mobility within them to allow the droplets to rearrange and coalesce, as seen at high temperature (Fig. ??Aii). Droplet fusion is driven by the surface tension and the viscosity of the droplets. These rheological properties determine if the final state of the coalescing process is complete, partial, or incomplete. If the surface tension dominates, as it does for simple liquids, the final state relaxes to a spherical shape. By this process, the surface curvature gradients are minimized to minimize surface energy. Otherwise, if the viscosity dominates, there would be slow to no relaxation of the condensates that results in non-spherical shapes.

As demonstrated in Fig. 1, we observe fusion events between two droplets (Fig. 1F). Prior studies have estimated the shape relaxation by fitting the boundary as it evolves from two spheres to one with a single ellipse with aspect ratio $l/w$, where $l$ is the semi-major axis and $w$ is the semi-minor axis (Fig. S7Ai). The aspect ratio shows an exponential relaxation over time with characteristic fusion time $\tau_e$ (Fig. S7Aii). A characteristic length scale of the process can be defined as: $L_e = \sqrt{w_0(l_0 - w_0)}$ (11, 12), where $l_0$ and $w_0$ are the semi-major and semi-minor axis lengths when the condensates are just in contact. For a simple fluid, $\tau_e$ is related to $L_e$ by $\tau_e \sim (\frac{\eta}{\gamma})L_e$, where $\frac{\eta}{\gamma}$ is the inverse capillary velocity and $\eta$ and $\gamma$ are the viscosity and the surface tension of the condensate, respectively(13). After fitting $\tau_e$ vs $L_e$ for MAP65 droplets with a linear equation $y = mx + c$, we find $\frac{\eta}{\gamma} \sim 0.38\ s/\mu m$ (Fig. S7Aiii). At a maturation time of 4.5 hours, we observed no change in aspect ratio in the coalescence process is arrested because of aging of the droplets (Fig. S7B).

Using the inverse capillary velocity and this viscosity deduced from FRAP measurements, we can estimate the surface tension of the condensates. In the first hour after the mix, the condensate surface tension is $\gamma \sim 12.3\ \mu N/m$. This surface tension value is reasonable compared to prior measurements for phase separated proteins, which vary from $\gamma \sim 0.1\ \mu N/m$ to $\gamma \sim 1\ mN/m$ (11, 14, 15). For comparison, air-water surface tension at 21.5 $^oC$ is, $\gamma \sim 72.75\ mN/m$, almost 6000 times larger than the estimated MAP65 surface tension value.

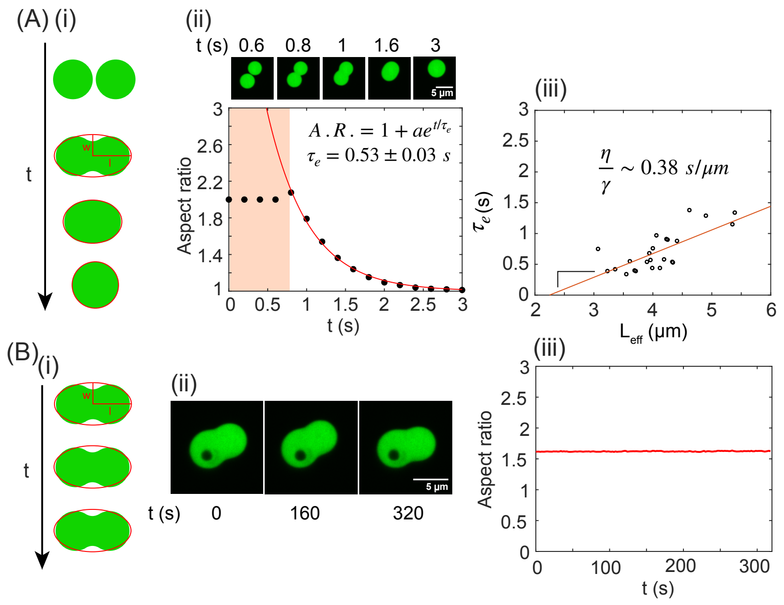


**Fig. S7.** Droplet merging quantification. (A) Fusion events at maturation time of one hour after mixing. (i) Cartoon schematic of the fusion event of two condensates over time. Time zero is when droplets first make contact, and an ellipse with semi-major axis $l$ and semi-minor axis $w$ is fit to outline the droplets over time. (ii) Image sequence of merging droplets showing time points t = 0.6 sec, 0.8 sec, 1 sec, 1.6 sec, and 3 sec. The scale bar is $5\ \mu m$. The aspect ratio from the displayed merging event, $l/w$, is plotted over time and shows an exponential decay. The fit to the data yields the characteristic fusion timescale, $\tau_e = 0.53 \pm 0.03$ s. (iii) The $\tau_e$ from several fusion events is plotted against the characteristic length scale, $L_e$. The linear slope represents the inverse capillary velocity $\eta/\gamma \sim 0.38\ s/\mu m$ (N = 26). (B) Characterization of fusion events at 4.5 hours of maturation time. (i) Cartoon schematic of the arrested fusion event. (ii) Image sequence of arrested droplets at time points t = 0 sec, 160 sec, and 320 sec. The dark dot is a pocket of buffer trapped region inside the condensate, and not due to photobleaching. This also demonstrates the gel-like nature of the droplets. (iii) The aspect ratio plotted against time stays uniform over longer time scale.

### References


1. KT Stanhope, JL Ross, *Microtubules, MAPs, and motor patterns.* (Elsevier Ltd) Vol. 128, pp. 23–38 (2015).
2. A Tulin, S McClerklin, Y Huang, R Dixit, Single-molecule analysis of the microtubule cross-linking protein MAP65-1 reveals a molecular mechanism for contact-angle-dependent microtubule bundling. *Biophys. J.* **102**, 802–809 (2012).
3. R Subramanian, et al., Insights into antiparallel microtubule crosslinking by PRC1, a conserved nonmotor microtubule binding protein. *Cell* **142**, 433–443 (2010).
4. B Edozie, et al., Self-organization of spindle-like microtubule structures. *Soft Matter* **15**, 4797–4807 (2019).
5. S Sahu, L Herbst, R Quinn, JL Ross, Crowder and surface effects on self-organization of microtubules. *Phys. Rev. E* **103**, 1–20 (2021).
6. R Dixit, JL Ross, *Studying plus-end tracking at single molecule resolution using TIRF microscopy.* (Elsevier) Vol. 95, First edit edition, pp. 543–554 (2010).
7. I Alshareedah, T Kaur, PR Banerjee, *Methods for characterizing the material properties of biomolecular condensates.* (Elsevier Inc.) Vol. 646, 1 edition, pp. 143–183 (2021).
8. CA Day, LJ Kraft, M Kang, AK Kenworthy, Analysis of protein and lipid dynamics using confocal fluorescence recovery after photobleaching (FRAP). *Curr. Protoc. Cytom.* (2012).
9. DM Mitrea, et al., Methods for Physical Characterization of Phase-Separated Bodies and Membrane-less Organelles. *J. Mol. Biol.* **430**, 4773–4805 (2018).
10. L Hubatsch, et al., Quantitative theory for the diffusive dynamics of liquid condensates. *eLife* **10**, 1–21 (2021).
11. S Elbaum-Garfinkle, et al., The disordered P granule protein LAF-1 drives phase separation into droplets with tunable viscosity and dynamics. *Proc. Natl. Acad. Sci. United States Am.* **112**, 7189–7194 (2015).
12. CP Brangwynne, TJ Mitchison, AA Hyman, Active liquid-like behavior of nucleoli determines their size and shape in Xenopus laevis oocytes. *Proc. Natl. Acad. Sci. United States Am.* **108**, 4334–4339 (2011).
13. DGAL Aarts, M Schmidt, HNW Lekkerkerker, Direct Visual Observation of Thermal Capillary Waves. *Science* **304**, 847–850 (2004).
14. CP Brangwynne, et al., Germline P granules are liquid droplets that localize by controlled dissolution/condensation. *Science* **324**, 1729–1732 (2009).
15. H Wang, FM Kelley, D Milovanovic, BS Schuster, Z Shi, Surface tension and viscosity of protein condensates quantified by micropipette aspiration. *Biophys. Reports* **1**, 100011 (2021).
